# Evolutionary history of sexual differentiation mechanism in insects

**DOI:** 10.1101/2021.11.30.470672

**Authors:** Yasuhiko Chikami, Miki Okuno, Atsushi Toyoda, Takehiko Itoh, Teruyuki Niimi

## Abstract

Gain of alternative splicing gives rise to functional diversity in proteins and underlies the complexity and diversity of biological aspects. However, it is still not fully understood how alternatively spliced genes develop the functional novelty. To this end, we infer the evolutionary history of the *doublesex* gene, the key transcriptional factor in the sexual differentiation of arthropods. *doublesex* is controlled by sex-specific splicing and promotes both male and female differentiation in some holometabolan insects. In contrast, *doublesex* promotes only male differentiation in some hemimetabolan insects. Here, we investigate ancestral states of *doublesex* using *Thermobia domestica* belonging to Zygentoma, the sister group of winged insects. We find that *doublesex* of *T. domestica* expresses sex-specific isoforms but is only necessary for male differentiation of sexual morphology. This result ensures the hypothesis that *doublesex* was initially only used to promote male differentiation during insect evolution. However, *T. domestica doublesex* has a short female-specific region and upregulates the expression of *vitellogenin* homologs in females, suggesting that *doublesex* may have already controlled some aspects of feminization in the common ancestor of winged insects. Reconstruction of the ancestral sequence and prediction of the protein structure show that the female-specific isoform of *doublesex* has a long C-terminal disordered region in holometabolan insects, but not in non-holometabolan species. We propose that *doublesex* acquired a female-specific isoform and then underwent a change in the protein motif structure, which became essential for female differentiation in sexual dimorphisms.

## Introduction

Sexual reproduction is widely used for transmitting genetic information from one to the next generation in Metazoa. For reproductive success, animals evolved diverse sex differences, i.e., sexual dimorphism, in morphology (Darwin 1871; Geddes and Thomson 1889) which underlie eco-evolutionary dynamics such as extinction rate and interspecific interaction (Fryxell et al. 2019). In the last three decades, the genetic pathways that create sex and sexual dimorphism have been elucidated in many animal species. Surprisingly, despite having a single origin (Beukeboom and Perrin 2014), these pathways have undergone extensive changes during animal evolution (Wilkins 1995; Bachtrog et al. 2014; Bopp et al. 2014; Herpin and Schartl 2015).

The diversity has been attributed to differences in the composition of the regulatory cascades. For example, in eutherians such as mice and humans, the master regulator of sex is *Sex-determining region Y* (*Sry*), a member of the High Mobility Group (HMG)-box transcriptional factor family (Gubbay et al. 1990; Sinclair et al. 1990; Koopman et al. 1991; Miyawaki et al. 2020), while *DM domain gene on the Y chromosome* (*dmy*) of the *doublesex* and *mab-3* related transcriptional factor (DMRT) family is the master sex-determining regulator in the medaka fish (Matsuda et al. 2002; Nanda et al. 2002). Diversification of the pathway governing sex determination/differentiation are largely based on differences in their gene repertoires (e.g., Hasselmann et al. 2008; Hattori et al. 2010; Sato et al. 2010; Takehana et al. 2014). In contrast, it has recently been discovered that the mechanisms of sexual differentiation in Pterygota, i.e., winged insects, differ in outputs of the gene cascade, e.g., the promotion masculinization or feminization.

Sexually dimorphic morphology in Pterygota is formed during postembryonic development. *doublesex* (*dsx*), a member of the DMRT family, acts as a global regulator at the bottom of the cascade to govern over sex differentiation (Kopp, 2012; Verhulst and van de Zande, 2015). In many pterygote insects studied, *dsx* is controlled by sex-specific splicing. In Diptera, Coleoptera, and Lepidoptera, sex-specific Dsx protein variants are essential for promoting either male or female differentiation in sexual dimorphism (e.g., Hildreth 1965; Burtis and Baker 1989; Ohbayashi et al. 2001; Kijimoto et al. 2012; Ito et al. 2013; Shukla and Palli 2012; Gotoh et al. 2016; Xu et al. 2017). For example, in the fruit fly *Drosophila melanogaster, dsx* is required to realize sex differences in external genitalia and foreleg bristle rows, while *dsx* mutants show an intersexual phenotype in these traits because both male and female differentiation are inhibited (Hildreth and Lucchesi 1963; Hildreth 1965). However, in the sawfly *Athalia rosae* (Mine et al. 2017, 2021), the silverleaf whitefly *Bemisia tabaci* (Guo et al. 2018), the brown planthopper *Nilaparvata lugens* (Zhuo et al. 2018), the German cockroach *Blattella germanica* (Wexler et al. 2019), and the damselfly *Ischnura senegalensis* (Takahashi et al. 2019, 2021), *dsx* has sex-specific isoforms and is responsible for male differentiation of morphological traits during postembryonic development, but not needed for female differentiation. Thus, despite expressing sex-specific isoforms, *dsx*’s role in sexual differentiation in Pterygota is different, as it controls both male and female differentiation or only male differentiation for sexual morphogenesis.

*dsx* in crustaceans and arachnids is reported to be highly expressed in males without sex-specific isoforms (Kato et al. 2011; Pomerantz et al. 2015; Li et al. 2018; Panara et al. 2019). Accordingly, *dsx* is only required for male differentiation of morphological traits in the water flea *Daphnia magna*. Wexler et al. (2019) proposed a stepwise evolution in which *dsx* had acquired sex-specific isoforms and later had become essential for female differentiation. However, roles of *dsx* are more diverse than expected. In Hymenoptera, *dsx* is involved in female differentiation of reproductive organs in the honeybee *Apis mellifera* (Roth et al. 2019), while *dsx* is non-essential for female differentiation in the sawfly *At. rosae* (Mine et al. 2017, 2021). In the milkweed bug *Oncopeltus fasciatus*, *dsx* is involved in both female and male differentiation of the genital organs (Just et al. 2021). *dsx* in *Be. tabaci* positively regulates the expression of a yolk precursor gene *vitellogenin* in females while it is not essential for female morphology (Guo et al. 2018), implying that *dsx* has different functionality for morphogenesis and otherwise in females. Overall, estimating the evolutionary history of *dsx* in Pterygota is still a challenging task. Also, it is unclear what factors led to the feminizing roles of *dsx* (Hopkins and Kopp 2021).

The phylogenetic distance between crustaceans and Pterygota and the lack of information about outgroups more closely related to Pterygota may be the reason for the gap of understanding of how *dsx* evolved from a monofunctional to a bifunctional regulator in arthropods. In an attempt to close this gap, we decided to include the firebrat *Thermobia domestica* (Zygentoma) in our analysis of *dsx*. Zygentoma is the sister group of Pterygota (Misof et al. 2014), does not copulate, and displays simple sexual dimorphisms, i.e., non-aedeagus male penises and female ovipositors (Kristensen 1975; Matsuda 1976; Emeljanov 2014; Beutel et al. 2017; Boudinot 2018). Otherwise, there is little difference in morphology between females and males, as Darwin (1871: 348) noted, “The sexes do not differ.” These features suggest that the level of sex differentiation in this species is very simple and likely to be ancestral. Thus, Zygentoma presents an ideal model for investigating the ancestral state of *dsx* in Pterygota. In this study, we investigated *dsx* in *T. domestica* and analyzed its functions in sexual differentiation. Also, we carried out the phylogenetic analysis, ancestral sequence reconstruction, and protein structure prediction to infer the evolutionary history of *dsx*.

## Results and Discussion

### Molecular evolution of *dsx* homologs and gene duplication of *dsx* in insects

Five *doublesex* (*dsx*) homologs were found in the transcriptome database of the firebrat *Thermobia domestica*. To identify which of them corresponds to the *dsx* ortholog in *T. domestica*, we compared these to *dsx* homologs found in transcriptome/genome/protein databases of various arthropods and vertebrates (supplementary table 1) and performed molecular phylogenetic analyses based on the amino acid sequences of their DNA-binding domains. As a result, the pancrustacean *dsx* was grouped into a distinct clade from the other DMRT family genes (fig. 1A). Within this clade, four subclades were recognized: Insect Dsx Clade 1, Insect Dsx Clade 2, Entognatha Dsx Clade, and Crustacea Dsx Clade. Insect Dsx Clade 1 contained the *dsx* found in Pterygota including *Drosophila melanogaster*. This clade also contained one of the 5 *dsx* homologs of *T. domestica*. We consider it likely that this *dsx* homolog is the corresponding ortholog of *T. domestica*. This Entognatha Dsx Clade also contained *dsx* from a springtail (Collembola) and a dipluran insect (Diplura). The Crustacea Dsx Clade contained *dsx* from branchiopods, including daphnids.

**FIG. 1.**
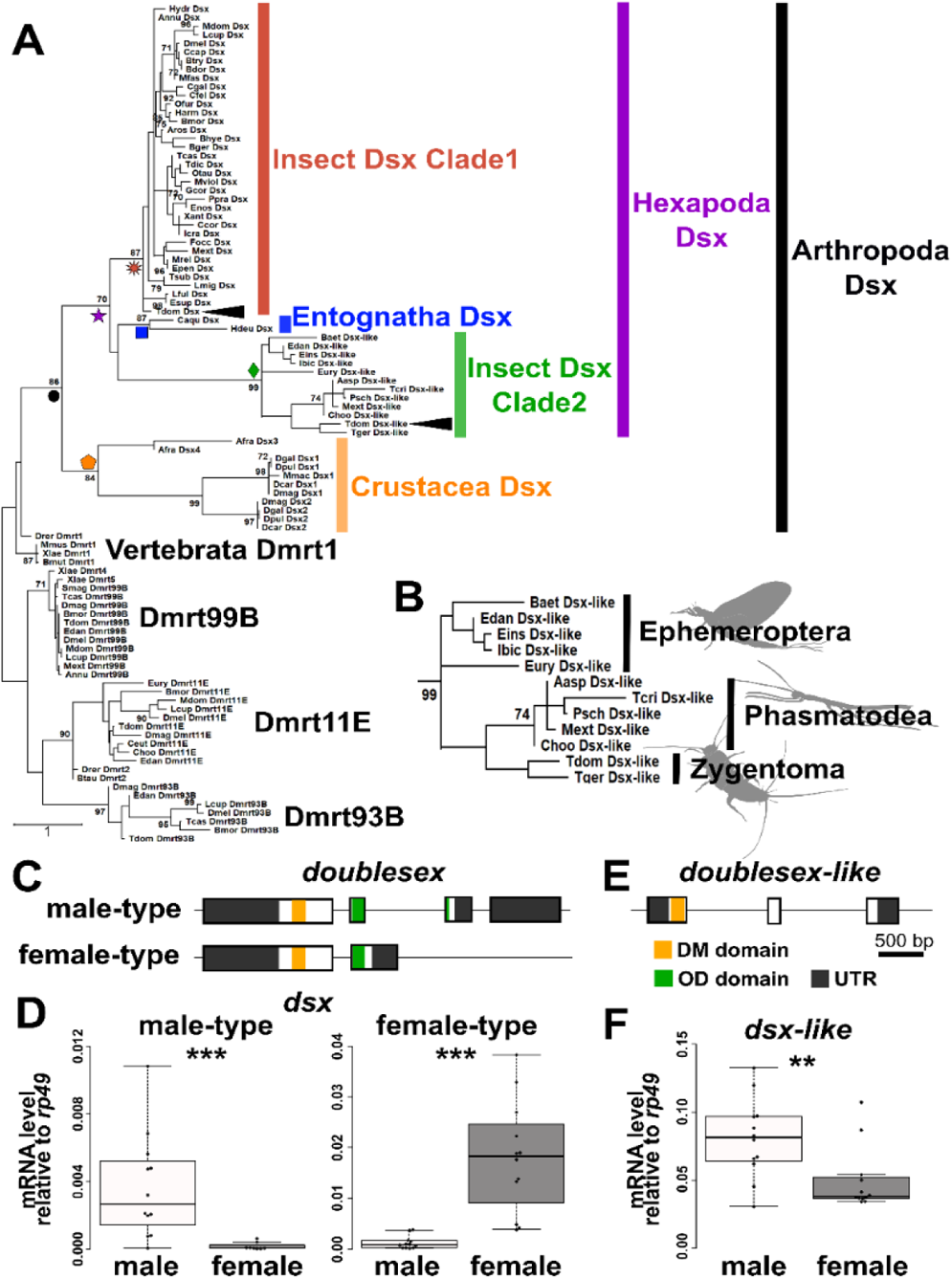
Molecular phylogeny and structural features of Doublesex in Arthropoda and Vertebrata. (*A*) Molecular phylogeny of Doublesex and Mab-3 related transcriptional factors (DMRT). The phylogenetic analysis was based on amino acid sequences of the DNA binding domain (DM domain) of DMRT family and was performed by the MEGA X after the multiple sequence alignment using the MAFFT software. The maximum-likelihood method was applied. 97 operational taxonomic units (OTUs) used for the phylogenetic analysis are listed in supplementary table 1. (B) Enlarged view of insect Dsx Clade2 (*dsx-like* clade). The numerical value on each node is the bootstrap supporting value. Bootstrap values < 70 are not shown. The node of each clade is indicated by colored shapes: black circle, Arthropoda Dsx; orange pentagon, Crustacea Dsx; purple star, Hexapoda Dsx; red sunburst, Insect Dsx Clade1; green diamond, Insect Dsx Clade2; blue square, Entognatha Dsx. (C) Exon-intron structures of *dsx* in *Thermobia domestica*. The upper and lower schematic images show the gene structure of *dsx* male-type and female-type, respectively. (D) Expression level of *dsx* in males and females of *T. domestica*. (E) Exon-intron structures of *dsx-like* of *T. domestica*. (F) Expression level of *dsx-like* in males and females. The exon-intron structure is determined by mapping the mRNA sequence of each gene to the genome of *T. domestica*. The expression level (D and F) was measured by the RT-qPCR of *dsx* and *dsx-like* in the adult fat body and is indicated as the relative values to the expression of the reference gene, *ribosomal protein 49* (*rp49*). Each plot indicates the mRNA expression level of each individual. Total *N* = 20 (*dsx* male-type), 23 (*dsx* female-type), and 24 (*dsx-like*). Results of Brunner–Munzel tests are indicated by asterisks: \*\**P*<0.01; \*\*\**P*<0.001 and are described in supplementary table 2.

We found that several species of Zygentoma, Ephemeroptera (mayflies), and Phasmatodea (stick insects) contain a *dsx-like* homolog of the Insect Dsx Clade 2 (fig. 1A, B) as well as the dsx ortholog of the Insect Dsx Clade 1. These findings indicate that *dsx* was duplicated before the divergence of Zygentoma and that the two paralogs retained from the divergence of the pterygote insects until at least the divergence of Eumetabola (= Hemiptera + Thysanoptera + Psocodea + Holometabola). The molecular evolution of *dsx* has been inferred from *dsx* of some pterygote insects, mainly holometabolan insects (Wexler et al. 2014; Mawaribuchi et al. 2019), while the presence of a *dsx-like* gene may have been overlooked in their analyses. Here, we report that the genome of *T. domestica* also contains both *dsx* and *dsx-like*, reflecting the presumed ancestral state in Pterygota in terms of gene copy number of *dsx*. Gene duplication generally leads to neo-/sub-functionalization to allow functional diversification (c.f., Taylor and Raes 2004). In this study, we analyzed the expression profiles and functions of *dsx* as well as *dsx-like* in *T. domestica*.

### Sex-specific splicing of *dsx* in *Thermobia domestica*

Splicing of *dsx* produces gives rise to sex-specific isoforms in all pterygote insects studied thus far, with the exception of the termite *Reticulitermes speratus* (Miyazaki et al. 2021), the silverleaf whitefly *Bemisia tabaci* (Guo et al. 2018), and the body louse *Pediculus humanus* (Wexler et al. 2019), suggesting that sex-specific splicing regulation of *dsx* was acquired before the divergence of Pterygota. To examine this hypothesis, we investigated the expression profile of *dsx* and *dsx-like* of *T. domestica*. Full-length mRNA sequences of *dsx-like* and *dsx* in *T. domestica* were determined by the RNA-seq and rapid amplification of cDNA ends (RACE) methods. Then, we investigated the gene structures to map the mRNA sequences to our genome database. *dsx* consists of five exons with two isoforms (fig. 1C): a long one (951 bp) and a short one (756 bp). RT-qPCR analysis showed that the long isoform and the short isoform were highly expressed in males and females, respectively (fig. 1D; Brunner-Munzel test, *P* = 1.75×10^−6^ and 2.20×10^−16^ in the long and the short isoforms). This fact indicates that *dsx* is controlled by sex-specific splicing. We refer to the male-biased isoform as *dsx* male-type and the female-biased isoform as *dsx* female-type. The *dsx-like* is expressed about two-fold higher in males than in females (Brunner-Munzel test, *P*=0.00924) and has three exons but no sex-specific isoform (fig. 1E, F), showing that *dsx-like* is not regulated by sex-specific splicing.

Our results give the further support that sex-specific splicing of *dsx* already existed in the common ancestor of Pterygota and Zygentoma (= Dicondylia), which diverged ~421 million years ago (Ma). Misof et al. (2014) estimated that the common ancestor of *Daphnia* and hexapods occurred at ~508 Ma. Therefore, *dsx* sex-specific splicing regulation is an ancient feature of insects that was acquired between 508 and 421 Ma and has been conserved for ~400 million years in each taxon of Dicondylia.

### Function of *dsx* for internal reproductive system and body size in *T. domestica*

Deciphering the role of *dsx* of *T. domestica* is essential for inferring the ancestral roles of *dsx* in pterygote insects. Hence, we conducted a functional analysis of not only *dsx* but also *dsx-like* since this paralog might also play a role in sexual differentiation.

To this end, we silenced *dsx* and *dsx-like* by RNA interference (RNAi). We quantified the expression of *dsx* and *dsx-like* in fat bodies of RNAi individuals by reverse transcriptional quantitative polymerase chain reaction (RT-qPCR). *dsx* silencing in females and *dsx-like* silencing in both sexes showed significantly decreased expression of each target genes compared to their expression in *enhanced green fluorescent protein* (*egfp*) RNAi controls (Brunner-Munzel test, *P* = 0.0265 in female *dsx*, 4.40×10^−16^ in male and female *dsx-like*; supplementary fig. 1A; supplementary table 2). The *dsx* RNAi males did not show a significant effect on *dsx* expression. Since it was suspected that outliers affected this result, we tested for outliers in *dsx* RNAi males and found one outlier (supplementary table 3). The reanalysis removing the outlier showed that *dsx* expression was significantly decreased in *dsx* RNAi males (Brunner-Munzel test, *P* = 0.00545; supplementary fig. 1B; supplementary table 2). Therefore, we concluded that *dsx* and *dsx-like* dsRNAs can knock down each target gene. Also, *dsx* RNAi had no effect on *dsx-like* expression and vice versa.

We also performed a double knockdown of *dsx* and *dsx-like*, to address the possibility that *dsx* and *dsx-like* are redundant. We specifically examined the effects of silencing on sexual dimorphism such as body size (fig. 2A) and reproductive systems. The body size, as measured by the pronotum width, was not affected in either knockdown group (fig. 2B; supplementary table 4, 5). In the gonads, the *dsx* RNAi, *dsx-like* RNAi, and double knockdown did not show any histological differences in testes, ovaries, and gametogenesis from the controls (fig. 2C; Supplementary Material online). On the other hand, in the *dsx* knockdown group (*dsx* alone or *dsx* and *dsx-like*), the male seminal vesicle, which is a sperm storage organ and normally has a bean pod shape, became rounded (fig. 2D). The number of sperm in the seminal vesicles of *dsx* RNAi males was lower than in the control group (fig. 2E; supplementary table 5; generalized linear model, *P* = 0.00487). Silencing of dsx or dsx-like or both did not affect normal differentiation of the female reproductive systems including the spermatheca (fig. 2F; supplementary fig. 2; Supplementary Material online). Also, there was no effect on the number of oocytes in any treatments (fig. 2G; supplementary table 4).

**FIG. 2.**
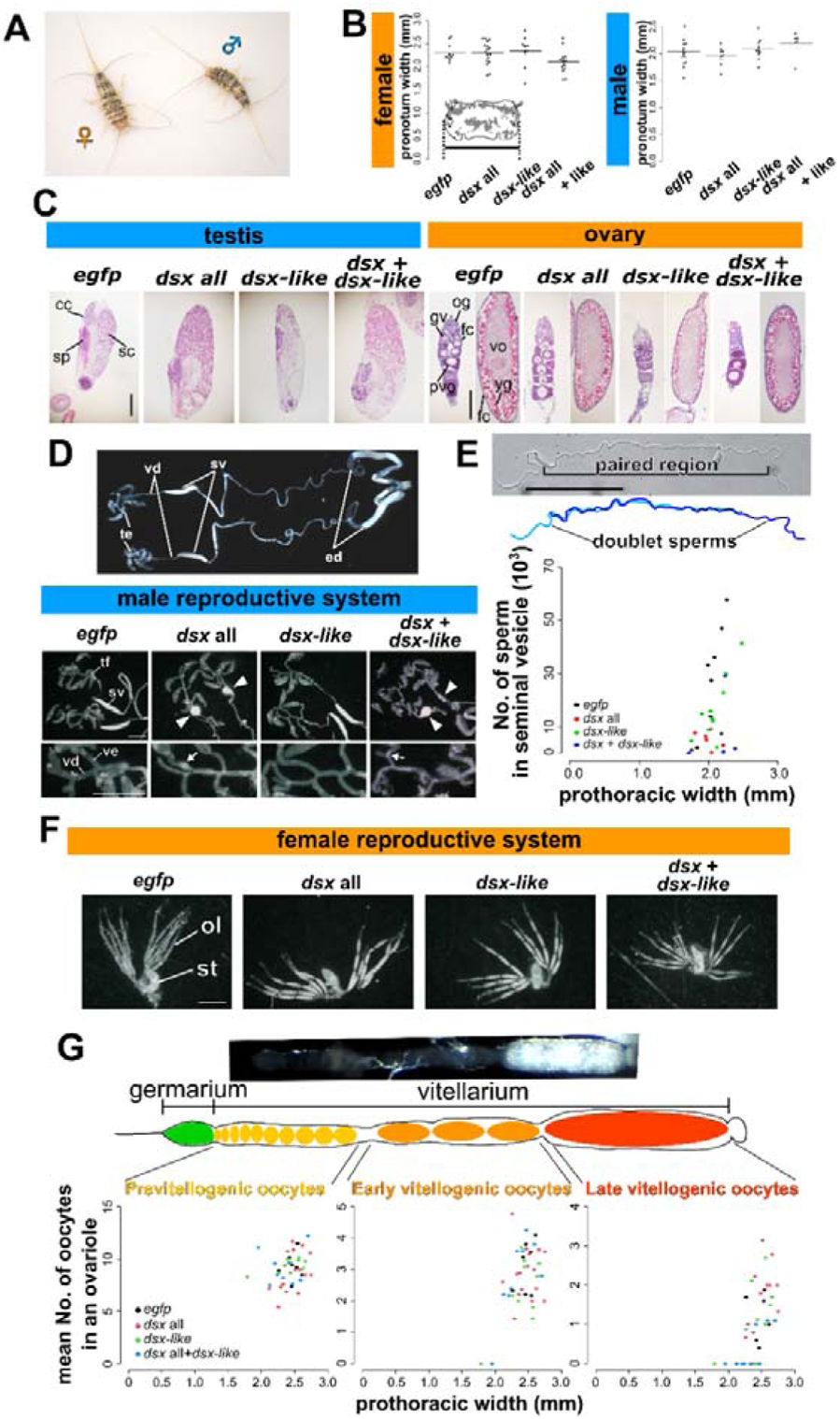
Function of *doublesex* and *doublesex-like* for body size, internal reproductive system, and gametogenesis of *Thermobia domestica*. (*A*) A pair of *T. domestica*. The female looks much the same as the male. (*B*) Body size of RNAi treatment groups. The pronotum (prothoracic tergum) width was used for the index of the body size. The graph shows mean ± SE (standard error). The results of the generalized linear model (GLM) analysis show in supplementary table 4 (female) and 5(male). Any significant effect can be detected in the RNAi treatments. Total *N* = 49 in females and 36 in males. (*C*) Histology of gonads in the RNAi groups. Paraffin. Hematoxylin-Eosin staining. In images of the ovary, the left and right panel in each treatment show germarium/previtellogenesis and vitellogenesis, respectively. (*D*) Effects of RNAi on male internal reproductive system. The upper photo shows the gross morphology of the reproductive systems in the non-treated male. The lower photos demonstrate the morphology of the RNAi males. The arrowheads show the rounded seminal vesicle. The lowest photos focused on the vas efferens. The arrows show the clogged sperm in the vas efferens. (*E*) Sperm of RNAi males. The upper photo and figure are sperm morphology in the non-treated male. The sperm forms doublet in the seminal vesicle. The lower figure shows the sperm number of the RNAi males. The results of the GLM analysis show in supplementary table 5. The significant effect was detected in the *dsx* RNAi treatment (*P* = 0.00487). Total *N* = 29. (*F*) Effects of RNAi on female internal reproductive system. (*G*) Effects of the RNAi on oocyte number. The upper photo shows the ovariole of the non-treated female. The lower figures exhibit the number of oocytes in the RNAi females along with the oogenetic stages. The results of the GLM analysis show in supplementary table 4. The number of the late vitellogenic oocytes was correlated with the pronotum width, although any significant effect can be detected in the RNAi treatments. Total *N* = 42 in each stage. In each panel, the *egfp*, *dsx* all, *dsx-lik*e and *dsx* + *dsx-like* indicates the *egfp* dsRNA injected group (control), *dsx* sex-common region dsRNA injected group, *dsx-like* dsRNA injected group, and both *dsx* sex-common region and *dsx-like* dsRNAs injected group, respectively. Each plot in (*B*), (*E*), and (*G*) indicates the value of each individual. cc, cystocyte; fc, follicle cell; gv, germinal vesicle; og, oogonia; ol, ovariole; pvo, previtellogenic oocyte; sc, spermatocyte; sp, sperm; st, spermatheca; sv, seminal vesicle; tf, testicular follicle; yg, yolk granule; ve, vas efferens; vd, vas deferens, vo, vitellogenic oocyte. Scales: 50 µm (*C*); 10 µm (*E*); 1000 µm (*D* and *F*).

The lack of effect of RNAi on the gonads may be due to the timing of the RNAi treatment, which was performed after gonadal differentiation. This is supported by a previous study (Klag 1977) suggesting that sex differences in gonads and germ cells are produced during embryogenesis. Embryonic RNAi is necessary to test this hypothesis, although, this experiment will be left to future studies, as our study focuses on the function of *dsx* during postembryonic development. The lack of effect of *dsx* on the body size of *T. domestica* is consistent with studies in *D. melanogaster* (Hildreth 1965; Rideout et al. 2015). The effect on internal reproductive systems other than the gonads are consistent with results in *Athalia rosae* (Mine et al. 2017, 2021) and *Blattella germanica* (Wexler et al. 2019).

### Function of *dsx* for morphology in genital organs of *T. domestica* and evolution of the function of *dsx* for sexual morphogenesis in insects

The sexually dimorphic morphology can be seen in the external genital organs, i.e., male penis and female ovipositor (fig. 3A). Males of *T. domestica* have unpaired small external genitalia on the abdominal segment IX. Females have an ovipositor consisted of two paired appendage-like structure on the abdominal segment VIII and IX. Males of the *dsx* knockdown groups (*dsx* only and both *dsx* and *dsx-like* RNAi) was transformed into two pairs of appendage-like structures resembling the female ovipositor (fig. 3B, C; Supplementary Material online). This effect was not observed in *dsx-like* RNAi males. Our results indicate that *dsx* is essential for male differentiation of morphological traits in *T. domestica*. In contrast to males, our analysis showed that none of the RNAi treatments affected female ovipositors at external morphological, tissue, or cellular levels (fig. 3D, E; supplementary fig. 3; Supplementary Material online). We then measured the length of the female ovipositor in the RNAi-treated groups to examine the involvement of *dsx* and *dsx-like* in the growth of female morphology. The results showed that *dsx* and *dsx-like* RNAi had no significant effect on ovipositor length (fig. 3F, G; supplementary table 4).

**FIG. 3.**
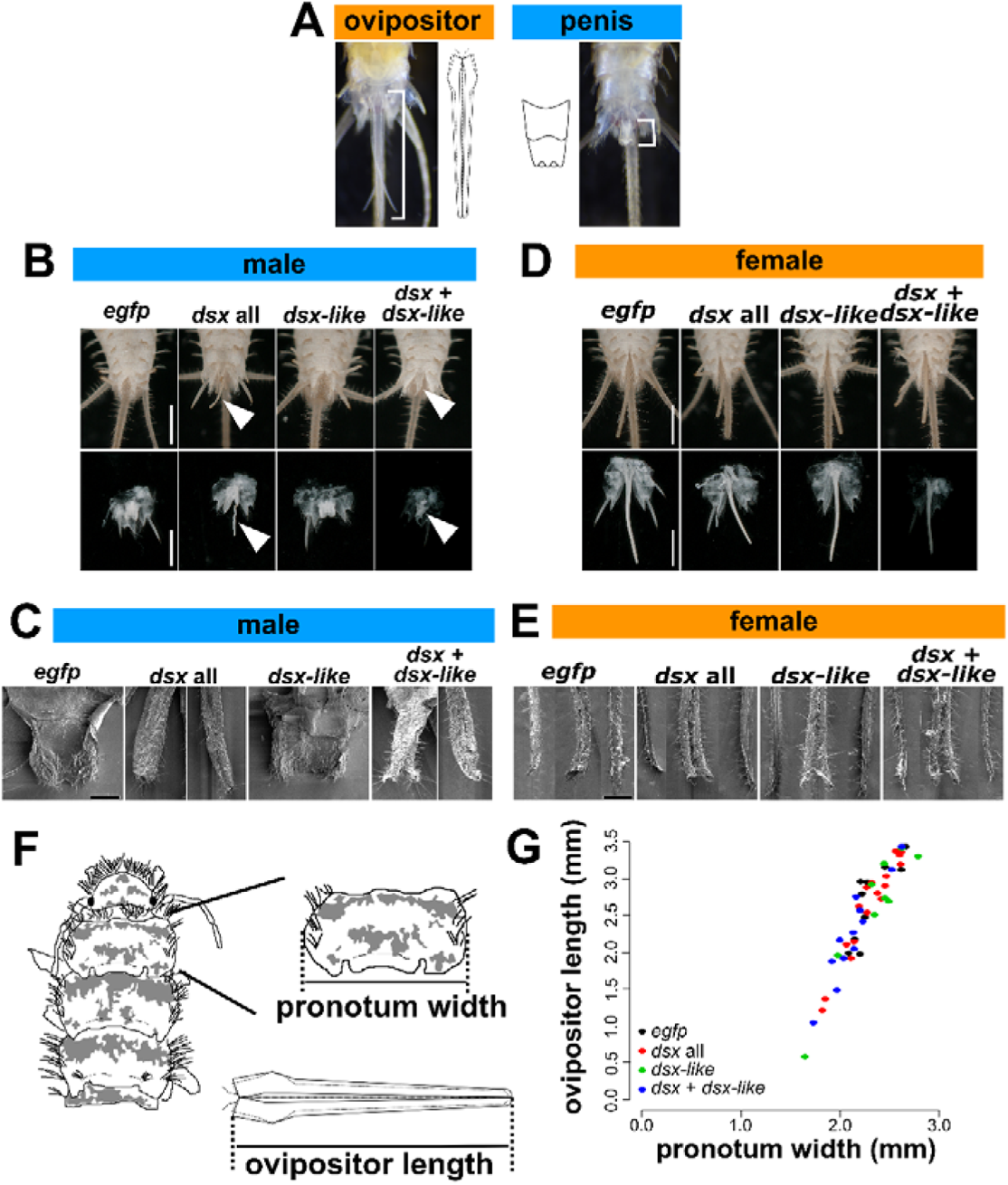
Function of *doublesex* and *doublesex-like* for genital organs in *Thermobia domestica*. (*A*) Sexually dimorphic traits of *T. domestica*. Females possess an ovipositor and males have a penis. (*B*) Effects of RNAi treatments on male penial structure. The upper images show the ventral side of the male abdomen. The lower images focus on the male penis. The arrowheads indicate the ovipositor-like structure in *dsx* or both *dsx* and *dsx-like* RNAi groups. (*C*) SEM images of male penial structure. In *dsx* and *dsx* + *dsx-like* RNAi, the two photos are merged into the one image. In these images, the left panels show the ovipositor valvula II (inner sheath) - like structure. The right panels exhibit the ovipositor valvula I (outer sheath)-like structure. The detail description can be referred in Supplementary Material online. (*D*) Effects of RNAi treatments on female ovipositor. The upper images show the ventral side of the female abdomen. The lower images focus on the female ovipositor. (*E*) SEM images of female ovipositor structure. In each image, the left and right panels show the valvula II and the middle one exhibits the valvula I. The results of the histological observation are in supplementary fig. 3. The detail description can be referred in Supplementary material online. (*F*) The schematic images of the measured parts. (*G*) Effects of RNAi treatments on growth of ovipositor. Each plot indicates the ovipositor length of each individual. The results of the generalized linear model analysis show in supplementary table 4. The ovipositor length was correlated with the prothoracic width (*P* = 2.00×10^−16^), although any significant effects can be seen in the RNAi treatments. Total *N* = 38. In each panel, the *egfp*, *dsx* all, *dsx-lik*e and *dsx* + *dsx-like* indicates the *egfp* dsRNA injected group (control), *dsx* sex-common region dsRNA injected group, *dsx-like* dsRNA injected group, and both *dsx* sex-common region and *dsx-like* dsRNAs injected group, respectively. Scales: 1 cm (*B* and *D*); 50 µm (*C* and *E*).

The lack of effect of *dsx* RNAi in females is due to that *dsx* is not essential for female differentiation of morphology during postembryonic development or that *dsx* knockdown is inefficient in the females. Compared to the knockdown efficiency of *dsx* in the males (~30% at median), that in the females is ~50% (supplementary table 2). However, given that half of the female individuals in the RT-qPCR analysis in the fat body had lower *dsx* expression than the minimum value of the control ones (supplementary fig. 1A), it can be assumed that *dsx* expression is suppressed in a certain number of females used in each analysis. In addition, *dsx* RNAi showed no effect on morphology in all 58 females (80 females including *dsx* and *dsx-like* double RNAi) analyzed in this study. Thus, it is reasonable to conclude that *dsx* is not essential for female differentiation of morphology during postembryonic development in *T. domestica*. Also, our results indicate that *dsx-like* is not essential for the sexual differentiation of morphology during postembryonic development in *T. domestica*. The knockdown of both *dsx* and *dsx-like* showed only the same effect as *dsx* RNAi alone. Thus, it is unlikely that *dsx-like* functions redundantly with *dsx*.

Sexual morphology, e.g., reproductive systems and genital organs, formed during postembryonic development is controlled by *dsx* in males but is *dsx*-independent in females of non-holometabolan insects such as *T. domestica* (Zygentoma: this study), *Bl. germanica* (Dictyoptera: Wexler et al. 2019) and the brown planthopper *Nilaparvata lugens* (Hemiptera: Zhuo et al. 2018). There could be the possibility of tissue-specific effects of *dsx* found in some holometabolan females (e.g., Ledón-Rettig et al. 2017). However, this possibility would be unlikely at least in these species since *dsx* was knocked down by systematic RNAi and was not reported to affect female morphology at this time. Based on these facts, we estimate that *dsx* may not be essential for female differentiation of morphology at the common ancestor of Dicondylia, ensuring the hypothesis proposed by Wexler et al. (2019).

To elucidate the timing of the acquisition of the role of *dsx* in female morphogenesis during postembryonic development, we must interpret the role of *dsx* in Hymenoptera, the basal clade of Holometabola. Studies in the honeybee *Apis mellifera* showed through genome editing that *dsx* controls female differentiation of the internal reproductive system under worker nutrition conditions (Roth et al. 2019). In the honeybee, sex differences in the gonads are established during embryogenesis (Lago et al. 2020). Thus, the male-like reproductive organ in *dsx* mutant females in Roth et al. (2019) would show an effect during embryogenesis, not during postembryonic development. We cannot conclude whether *dsx* is not essential for female morphogenesis in the honeybee, since the information on the roles of *dsx* in sexual morphology is limited to gonads and heads of the worker females. However, given that *dsx* does not affect heads in *Ap. mellifera* females (Roth et al. 2019), wings in the parasitoid wasp *Nasonia vitripennis* females (Wang et al. 2020), and sexual traits in *At. rosae* females (Mine et al. 2017, 2021), at this time, it is reasonable to infer that *dsx* was not essential for female morphogenesis during postembryonic development in the common ancestor of Hymenoptera. This interpretation and the essential roles of *dsx* for female development in the other holometabolan insects suggest that *dsx* became essential for feminization of morphology during postembryonic development at the common ancestor of holometabolan insects except for Hymenoptera (=Aparaglossata) emerging ~327 Ma.

### Cryptic role of *doublesex* for female-specific transcripts in *T. domestica* and its opposite role between sexes

*dsx* in *T. domestica* does not seem to have conflicting functions between sexes in postembryonic morphogenesis. On the other hand, other biological processes remain to be considered. We tested whether *dsx* contributes to the expression of *vitellogenin* (*vtg*), a yolk protein precursor gene that is highly expressed in animal females (Byrne et al. 1989; Hayward et al. 2010). Previous studies have shown that *vtg* in pterygote insects is controlled by *dsx* (e.g., Suzuki et al. 2003; Shukla and Palli 2012; Thongsaiklaing et al. 2018). Our RNA-seq analysis showed that three *vtg* homologs, i.e., *vtg1*, *vtg2*, and *vtg3,* were expressed female-specifically in the fat body in *T. domestica* (supplementary fig. 4; supplementary table 6). We analyzed the expression of *vtg* in the fat bodies of *dsx, dsx-like,* or both genes RNAi groups by RT-qPCR.

In *dsx* RNAi males, all *vtg* mRNAs were expressed 45–1530-fold higher than the controls (fig. 4A; supplementary table 2: Brunner-Munzel test, *P* = 2.87×10^−8^, 6.60×10^−16^ and 2.80×10^−4^ in *vtg1*, *vtg2*, and *vtg3*). *vtg1* and *vtg3* mRNAs were significantly up-regulated in *dsx-like* RNAi males compared to the controls (fig. 4A: Brunner-Munzel test, *P* = 0.0139 and 0.00497 in *vtg1* and *vtg3*). In both *dsx* and *dsx-like* RNAi males, the effect was similar to that in *dsx* RNAi males (fig. 4A: Brunner-Munzel test, *P* = 6.60×10^−16^, 0.0162, and 6.60×10^−16^ in *vtg1*, *vtg2*, and *vtg3*). Then, we found that the expression of all *vtg* genes was significantly reduced in *dsx* RNAi females (fig. 4B; supplementary table 2: Brunner-Munzel test, *P* = 0.0433, 0.00422, and 0.00623 in *vtg1*, *vtg2*, and *vtg3*). This reduction rate was approximately 0.2–0.4-fold. Furthermore, *vtg* expression was significantly reduced in *dsx-like* RNAi females (Brunner-Munzel test, *P* = 0.00256, 3.80×10^−6^, and 1.49×10^−5^ in *vtg1*, *vtg2*, and *vtg3*) and both *dsx* and *dsx-like* RNAi females (Brunner-Munzel test, *P* = 0.0305, 0.00892, and 0.0197 in *vtg1*, *vtg2*, and *vtg3*) (fig. 4B). These results show that *dsx* and *dsx-like* of *T. domestica* control *vtg* negatively in males and positively in females.

**FIG. 4.**
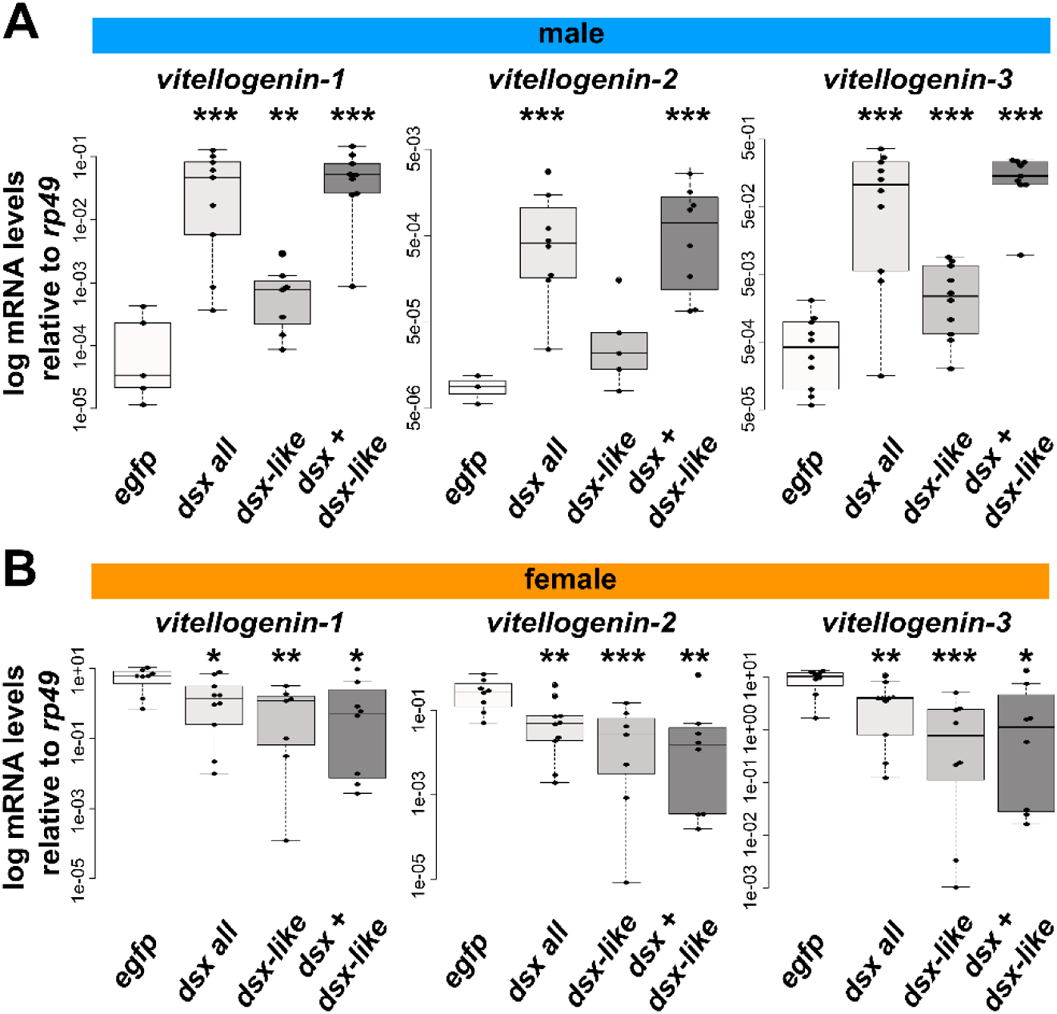
Function of *doublesex* for *vitellogenin* expression in *Thermobia domestica*. (*A*) *vitellogenin* expression level in RNAi males. (*B*) *vitellogenin* expression level in RNAi females. The mRNA expression levels were measured by the RT-qPCR analysis. The figures show the log-scale relative values of the expression level of three *vitellogenin* homologs to the reference gene, *ribosomal protein 49* (*rp49*). Each plot indicates the mRNA expression level of each individual. In each panel, the *egfp*, *dsx* all, *dsx-lik*e and *dsx* + *dsx-like* indicates the *egfp* dsRNA injected group (control), *dsx* sex-common region dsRNA injected group, *dsx-like* dsRNA injected group, and both *dsx* sex-common region and *dsx-like* dsRNAs injected group, respectively. The Brunner-Munzel test method were performed to statistically analyze the difference in mRNA expression level between the control and the *dsx* or *dsx-like* RNAi groups. The *P*-values were adjusted by the Holm’s method. \**P*<0.05, \*\**P*<0.001, \*\*\**P*<0.0001. *P* ≥ 0.05 is not shown. The statistical results were described in supplementary table 2. Total *N* = 30 (*vitellogenin-1*), 24 (*vitellogenin-2*) and 39 (*vitellogenin-3*) in males and 33 (*vitellogenin-1*), 33 (*vitellogenin-2*), and 34 (*vitellogenin-3*) in females.

Our results indicate that *dsx* has opposite roles between sexes, i.e., repressive in males and promotive in females, in *vtg* expression. *dsx-like* also has the opposite functions for *vtg* expression in males and females. It is unlikely that this result is due to *dsx-like* regulating *dsx* transcription, as *dsx-like* did not affect *dsx* expression (supplementary fig. 1A). A possible hypothesis is that *dsx-like* might regulate *vtg* expression as one of the co-regulators that bind *dsx* or other transcription factors.

We do not know whether *dsx* of *T. domestica* oppositely controls genes other than the *vtg* homologs between sexes since our analysis was limited to *vtg* homologs. However, the results from these genes indicates that the molecular function of *dsx* in this species includes the opposite function for some genes’ transcription in females and males. In *Be. tabaci*, *dsx* positively regulates *vtg* expression in females, even though it is not essential for female differentiation of morphological traits (Guo et al. 2018). *dsx* of this species does not negatively regulate *vtg* in males. Therefore, the functionality of *dsx* found in *T. domestica*, i.e., the opposing role in some genes’ expression between sexes and the function that are not essential for female morphogenesis, is a functionality that has not been reported in any insect or animal. This functionality indicates that even if *dsx* can oppositely function for some genes’ expression between sexes, it does not necessarily have opposite functions in morphogenesis between sexes. This difference in the functionality might be due to differences in genes under *dsx* control between morphogenesis and other aspects such as the yolk synthesis in females.

Genes under *dsx* control in males are *dsx*-free in females of *I. senegalensis* (Takahashi et al. 2021), *Bl. germanica* (Wexler et al. 2019; Pei et al. 2021), and *Ni. lugens* (Zhuo et al. 2018). It was thought that feminizing roles of *dsx* in morphogenesis and other biological processes may have appeared in the common ancestor of Aparaglossata (or Holometabola) as an entirely novel function, i.e., neofunctionalization. In contrast, the contribution of *dsx* to some genes’ expression in females of *T. domestica* (this study), *Be. tabaci* (Guo et al. 2018), *Ap. mellifera* (Velasque et al. 2018) and Aparaglossata raises the alternative hypothesis that the ability of *dsx* to be involved in female differentiation was already present in the common ancestor of Dicondylia and later became essential for the female morphogenesis in the common ancestor of Aparaglossata. In this evolutionary scenario, the role of *dsx* in the feminization of postembryonic morphogenesis in Aparaglossata could be due to extending its capability to control some genes in females, i.e., functional expansion. We cannot decide which of these hypotheses is appropriate, at this time. However, the latter scenario can well explain the presence of female-specific coding sequences of *dsx* and high expression of *dsx* female-type during postembryonic development, in non-aparaglossatan insects. The capability to regulate some female genes might be a “minor function” of *dsx* in non-holometabolan females predicted by Wexler et al. (2019).

### Evolution of C-terminus disordered region of *dsx* female-type

One of the puzzling problems is how *dsx* became recruited for female differentiation of morphological traits (cf., Hopkins and Kopp 2021). Here, we found that the C-terminal sequences including the oligomerization (OD) domain of the *dsx* female-type is much shorter in *T. domestica* (38 aa) than that in *D. melanogaster* (53 aa) (supplementary fig. 5). The OD domain is essential for female differentiation in *D. melanogaster*, as it physically binds to *dsx* itself, transcription factors, and co-activators (An and Wensink 1995; Erdman 1996; Ghosh et al. 2019; Romero-Pozuelo et al. 2019). Therefore, we hypothesized that the additive region found in *D. melanogaster* occurred at the common ancestor of Aparaglossata in which *dsx* became essential for female morphogenesis. To test this hypothesis, we obtained sequences of *dsx* female-type from 48 insect species based on the National Center for Biotechnology Information (NCBI) protein/transcriptome shotgun assembly database and previous studies (supplementary table 7) and reconstructed ancestral sequences of *dsx* female-type. Our ancestral sequence reconstruction revealed that the C-terminal 16-amino acid region of *dsx* female-type found in the common ancestor of Aparaglossata was absent in the common ancestor of the other taxon (fig. 5A; supplementary fig. 6; supplementary table 8). This motif is conserved within Aparaglossata in our dataset although moderate sequence diversification was observed (supplementary fig. 6). In our dataset, almost all sequences of this motif were not found in species in which *dsx* is not essential for the female differentiation of morphological traits during postembryonic development. Exceptionally, *dsx* of *At. rosae* had an amino acid sequence in the region corresponding to this motif, but our results of ancestral sequence reconstruction showed that this sequence was acquired in parallel with Aparaglossata.

**FIG. 5.**
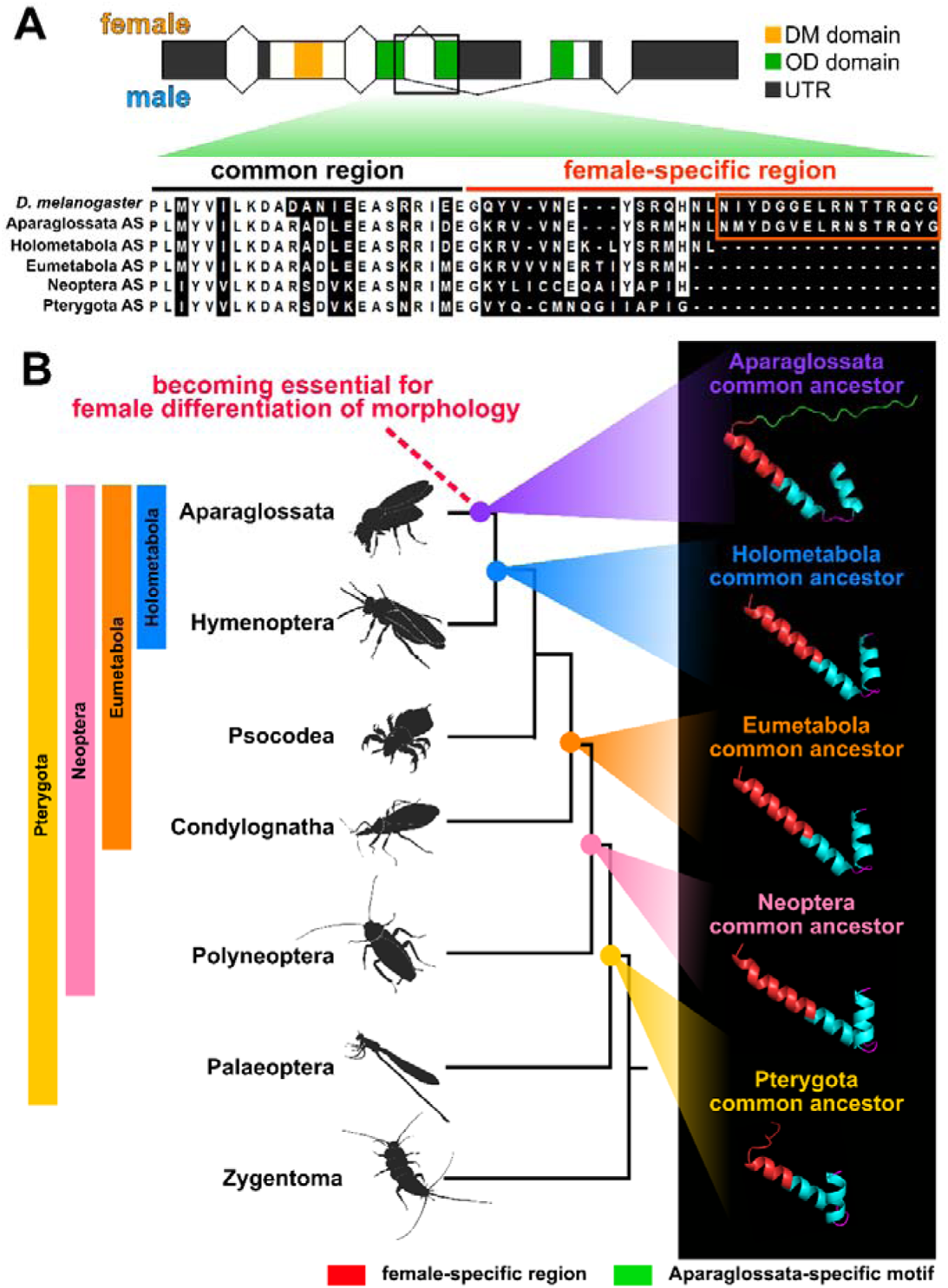
Evolution of C-terminal sequence of *doublesex* in insects. (*A*) Ancestral sequences (AS) of *dsx* in insects. The AS were reconstructed from 49 *dsx* proteins of insects by the maximum likelihood methods of the MEGA X. The information on the species and proteins used for the AS reconstruction is listed in supplementary table 7. The most probable sequences were applied. The results of the AS reconstruction are described in supplementary table 8. The upper scheme indicates the *dsx* gene structure of *D. melanogaster*. The lower image shows the result of the multiple sequence alignments (MSA) of *dsx* sequences by MAFFT. The oligomerization domain sequences at C-terminal side were used for the MSA. The white background in the MSA result indicates the conserved sites that share the residues in the 80% taxa. The Aparaglossata-specific motif is indicated by the orange frame. (*B*) Predicted protein structures of *dsx* female-type in common ancestors of insect taxa. The phylogenetic relationship is based on the topology of Misof et al. (2014). The 3D images in the right panel indicate the predicted structures of the OD domain including the female-specific region of *dsx*. The protein structures were predicted by the AlphaFold2-based algorism (ColabFold: Mirdita et al. 2021). The red region of the 3D image indicates the female-specific region. The green region shows the Aparaglossata-specific motif. The information on the evaluated values (predicted local distance difference test: plDDT) of the prediction is shown in the Material and Methods section and supplementary fig. 10.

The Aparaglossata-specific region is located in the distal (C-terminal) part of the female-specific region in *D. melanogaster*. This distal region is a disordered region, i.e., a mobile region that lacks a fixed structure, following an α-helix loop in the proximal (N-terminal side) region (Yang et al. 2008). To investigate whether the acquisition of the disordered region occurred in Aparaglossata, we predicted the protein structure of *dsx* female-type ancestral sequences of Pterygota, Neoptera, Eumetabola, Holometabola, and Aparaglossata. According to the Alphafold2 algorism-based structure prediction, the female-specific region of *dsx* in the common ancestor of Aparaglossata had a proximal α-helix loop structure, and a distal random coil indicating a disordered region (fig. 5B). This structure was similar to that of *D. melanogaster* determined by a crystal structural analysis (Yang et al. 2008). The proximal α-helix loop structure was also predicted in the common ancestors of taxon other than Aparaglossata. The random coil following the α-helix structure was predicted in all common ancestors, but its length was shorter than that of the common ancestor of Aparaglossata. It is essential to determine the structure via nuclear magnetic resonance or cryo-electron microscopy methods to conclude the details at the structural level, although, our theoretical predictions suggest that the disordered region following the α-helix structure in the female-specific region may have been extended in the common ancestor of Aparaglossata.

Our results suggest that both the extension of the disordered region following the α-helix loop in the female-specific region of *dsx* and the feminizing function of *dsx* for morphology occurred in the common ancestor of Aparaglossata. At present, the causality between these two events is uncertain, as we do not know which of the events appeared earlier. In general, disordered regions in transcription factors play essential roles in transcriptional activity through post-translational modifications and binding to co-activators and nucleic acids (Liu et al. 2006; Darling and Uversly 2018). Furthermore, Wang et al. (2019) showed that in the diamondback moth *Plutella xylostella,* when the Aparaglossata-specific motif is specifically broken by deletion or frameshift mutations using the CRISPR/Cas9 method, the female morphology is transformed into the intersexual phenotype. This result indicates that the Aparaglossata-specific motif is essential for female differentiation of morphology in *P. xylostella*. These facts suggest that the extension of the C-terminal region of *dsx* female-type may have been a key event associated with the acquisition of the female-differentiating roles of *dsx* in morphology during postembryonic development. This functional evolution of the “non-functional” isoform by the coding mutation is also consistent with an evolutionary process of alternative splicing isoforms theoretically predicted lacking empirical evidence (Keren et al. 2010).

### On the origin of outputs of the sexual differentiation mechanism

Recent findings in insects (e.g., Mine et al. 2017; Guo et al. 2018; Zhuo et al. 2018; Wexler et al. 2019; Takahashi et al. 2021), including this study, have shown that sexual differentiation mechanisms are diverse in their outputs as well as their gene repertoires. The diversity in the output is attributed to the functional diversity of a single gene, *dsx*, for sexual differentiation of morphogenesis during postembryonic development. The evolutionary origin of the diversity in the output is one of the enigmatic problems in sexual development. Information on the roles of *dsx* is limited to some traits in some species and cannot be available in many non-aparaglossatan species although functional analyses of *dsx* have been rapidly progressing using emerging model species. Unquestionably, comprehensive information on functions of *dsx* for sexually dimorphic morphology from wider taxa is essential for fully tracing the evolution of *dsx*. We propose, albeit premature, as one of the possibilities to be considered, the hypothesis by which *dsx* might have become essential for female differentiation in sexual morphology by expanding its cryptic feminizing role, i.e., functions for some female genes’ expression, in association with mutations in female-specific motifs (fig. 6). This scenario can explain how single genes acquire novel outputs of sexual development although our hypothesis does not prevent any other alternative hypothesis from being proposed.

**FIG. 6.**
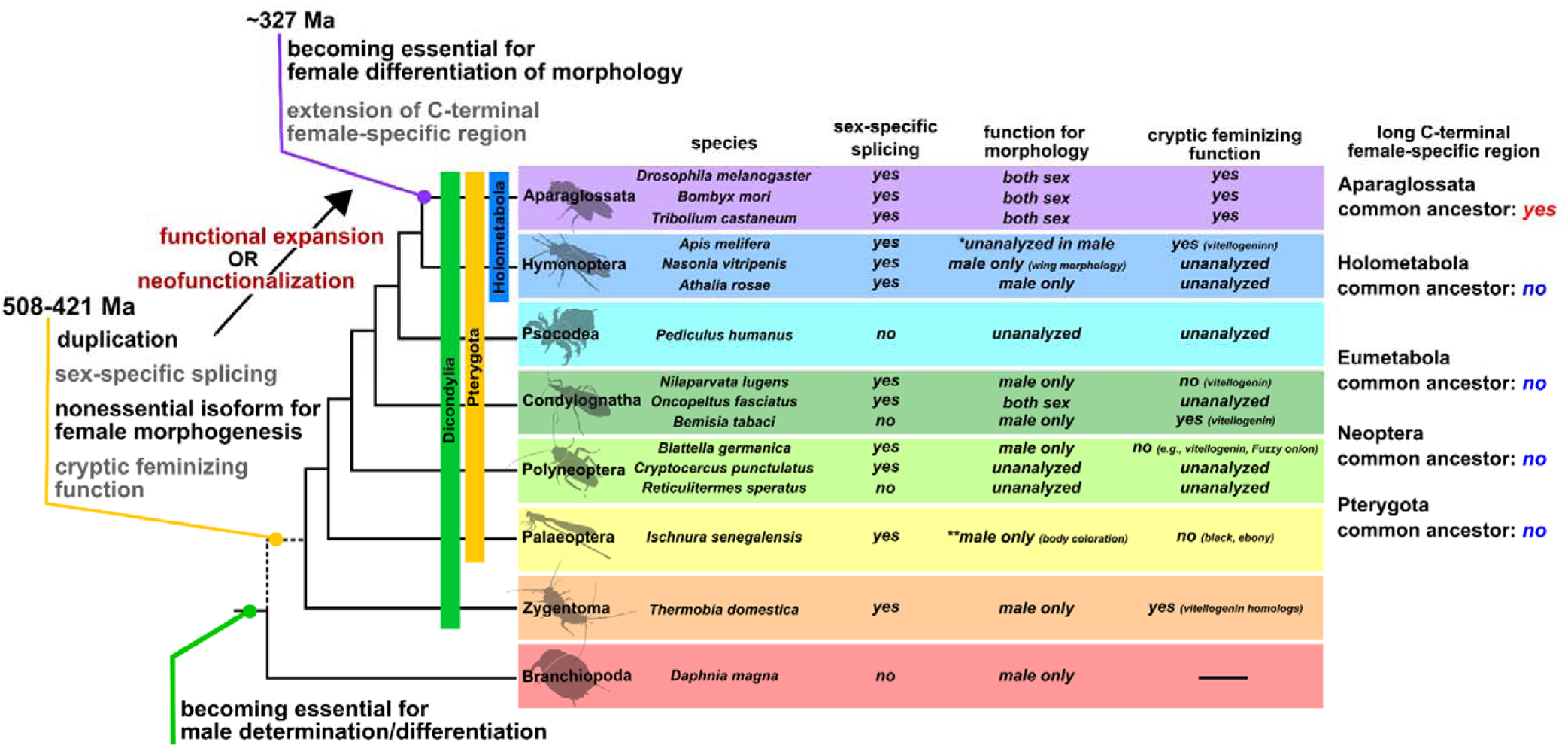
Schematic image of the evolutionary scenario of *doublesex* proposed in this study and the feature of *dsx* in insects. *dsx* in arthropods may have been initially involved in only male determination/differentiation based on findings in crustaceans and chelicerates (Kato et al. 2011; Pomerantz et al. 2015; Li et al. 2018; Panara et al. 2019). In our hypothesis, the female isoform of *dsx* may have been not essential for female differentiation of morphological traits at least from the common ancestor of Dicondylia to the common ancestor of Aparaglossata but might have contributed to the expression of some genes in females (“cryptic feminizing function” in the figure). This “seemingly non-functional” isoform might have become essential for female differentiation of morphological traits at the common ancestor of Aparaglossata emerging at ~327 million years ago (Ma) through extending its cryptic feminizing function (“functional expansion” in the figure) or, alternatively, through acquiring entirely novel function (“neofunctionalization” in the figure). The extension of the C-terminus amino acid sequences in the female-specific region might be involved in the functional expansion/neofunctionalization of *dsx*. The common ancestor between Branchiopoda and Hexapoda may have had the male-specific expressed *dsx*. It is estimated that the sex-specific splicing control and the gene duplication of *dsx* occurred from the common ancestor between Branchiopoda and Hexapoda emerging at ~508 Ma to the common ancestor of Dicondylia at ~421 Ma. The phylogenetic relationship and the divergence time refer to Misof et al. (2014). The dotted line in the phylogenetic relationship indicates that the taxa that occurred from the common ancestor between Branchiopoda and Dicondylia to the common ancestor of Dicondylia are omitted. Here, we also show information on the current knowledge of *dsx* features in insects and a branchiopod. In Aparaglossata, since there are many studies, we show only three representative species. The information was based on: Hildreth (1965), Bruce and Baker (1989) and Cloudh et al. (2014) in *Drosophila melanogaster* (Diptera), Ohbayashi et al. (2001), Suzuki et al. (2003), and Xu et al. (2017) in *Bombyx mori* (Lepidoptera), Shukla and Palli (2012) in *Tribolium castaneum* (Coleoptera), Roth et al. (2019) and Velasque et al. (2018) in *Apis mellifera* (Hymenoptera), Wang et al. (2020) in *Nasonia vitripenis* (Hymenoptera), Mine et al. (2017, 2021) in *Athalia rosae* (Hymenoptera), Wexler et al. (2019) in *Pediculus humanus* (Psocodea) and *Blattella germanica* (Dictyoptera), Zhuo et al. (2018) in *Nilaparvata lugens* (Hemiptera), Just et al. (2021) in *Oncopeltus fasciatus* (Hemiptera), Guo et al. (2018) in *Bemisia tabaci* (Hemiptera), Miyazaki et al. (2021) in the wood roach *Cryptocercus punctulatus* and *Reticulitermes speratus* (Dictyoptera), Takahashi et al. (2019, 2021) in *Ischnura senegalensis* (Odonata), this study in *Thermobia domestica* (Zygentoma), and Kato et al. (2011) in *Daphnia magna* (Branchiopoda). In Condylognatha, information on *dsx* in the blood-sucking bug *Rhodnius prolixus* is omitted. *R. prolixus* has sex-specific isoforms of *dsx* whose function has not been investigated (Wexler et al. 2014). The “unanalyzed” means the functional analyses of *dsx* have not been performed in the relevant species. Information on the roles of *dsx* of some species in female morphogenesis is limited to some body parts: e.g., body coloration in *I. senegalensis* (Takahashi et al. 2021), wing morphology in *Na. vitripenis* (Wang et al. 2020), and worker morphology in *Ap. mellifera* (Roth et al. 2019). The asterisk (*) in *Ap. mellifera* indicates that the functional analysis of *dsx* in males was not conducted although the gonad differentiation of female workers was affected by *dsx* knockouts (Roth et al. 2019; see Main text). The double-asterisk (**) in *I. senegalensis* shows that this species has polymorphic coloration in females, i.e., gynomorph (normal female color) and andromorph (male-like color) and that *dsx* is involved in the color formation of the males and andromorphic females but not gynomorphic females (see Takahashi et al. 2021), suggesting that *dsx* is not essential for the female color development. In our hypothesis, the essential roles of *dsx* for female development in *O. fasciatus* (Just et al. 2021) may have occurred in parallel with Aparaglossata.

The diversity of mechanisms that produce animal sex is a model case of developmental systemic drift (True and Haag 2001; Haag and True 2021). The functional diversity of a single gene and its evolutionary process has not been focused on in the context of developmental system drift to date due to its poor examples. Our evolutionary scenario may be one hypothesis explaining the origin of the system drift in the function of single genes. In this study, we have mainly discussed the functionality of *dsx* for sexual differentiation of morphology during postembryonic development. Therefore, it is unclear whether our conclusions and evolutionary scenarios apply to sexual behavior, including sex pheromone secretion and courtship, as well as sexual determination and gonadal differentiation during embryogenesis.

Detailed studies of sex differences at various levels across insect taxa will test our evolutionary scenario and will fully reconstruct the evolutionary history of *dsx* and sexual differentiation mechanisms.

## Materials and Methods

### Animals

The firebrat, *Thermobia domestica* (Packard 1873), was used as an emerging model for apterygote. *T. domestica* is one of the species belonging to Zygentoma (Lepismatidae). The insects were kept at 37°C in total darkness condition and fed with fish food (TetraFin Goldfish Flakes, Tetra GmbH, Melle, Germany) in our laboratory. Stock colonies were reared in plastic cases of 30 cm×40 cm or 18 cm × 25 cm in length. Eggs were collected from tissue paper in the case and incubated at 37°C. For examining the roles of *dsx* and *dsx-like* in the postembryonic morphogenesis, colonies of hatched nymphs were reared up to the fourth instar in a six-well plate and then transferred into 24-well plates to be kept individually. For examining the roles of *dsx* and *dsx-like* in *vitellogenin* expression, female and male insects were collected from the stock colony and transferred into the plates. For examining the function of *dsx* and *dsx-like* for sexual morphology and gametogenesis, we used firebrats from April to June, 2019, February to April, April to July, and September to December, 2020. For investigating the roles of *dsx* and *dsx-like* in the *vitellogenin* expression, firebrats were manipulated from June to July, 2020.

### Estimation of molt timing

Estimating the molt timing of insects is essential for the analysis of developmental processes and the functions of developmental regulatory genes. The timing of Hemi- or holometabolan insects can be estimated using morphological changes such as a wing growth. However, timing is hard to estimate in apterygote insects since they have little change in their morphology during postembryonic development. *T. domestica* forms scales in the fourth instar, and changes the number and length of its styli during the fourth to ninth instar under our breeding conditions. These features can be used to estimate molt timing, but it is difficult to apply these criteria to experiments using adults or a large number of nymphs. To resolve this problem, we used leg regeneration after autotomy and time-lapse imaging to estimate the molt timing of *T. domestica*. Autotomy occurs at the joint between the trochanter and femur in *T. domestica*. An autotomized leg regenerates after one molt (*Buck and Edwards, 1990*). For the RNAi analysis during postembryonic development, we amputated a right hindleg at the autotomic rift, using tweezers, and observed whether the leg had regenerated. This test enabled us to rapidly estimate the molt timing. For the RNA-seq and the RT-qPCR analysis, the time-lapse imaging was used to determine the precise time of molt. We build a time-lapse imaging system with a network camera system (SANYO, Tokyo, Japan) set in an incubator at 37°C (supplementary fig. 7A). Photos of insects in the 24-well plate were taken every five minutes. We created a time-lapse movie from the photos every 12 hours using ImageJ 1.52a (https://imagej.nih.gov/ij/) and observed whether the insects molted (supplementary fig. 7B).

### De novo genome assembly

A whole genome of *T. domestica* was sequenced to analyze the exon-intron structure of *dsx*. We selected an adult female of *T. domestica* from our stock colony and removed its alimentary canal. Genomic DNA was extracted from the sample using DNeasy Blood and Tissue Kit (QIAGEN K.K., Tokyo, Japan). A paired-end library was constructed from 1 µg of the DNA using TruSeq DNA PCR-Free LT Sample Prep kits (Illumina K.K., Tokyo, Japan) following the manufacturer’s instructions. The library was run on a sequencer (HiSeq 2500; Illumina K.K., Tokyo, Japan). We obtained 417 Gb of raw reads and assembled them using Platanus v1.2.4 assembler (*Kajitani et al. 2014*) after removal of the adapter sequences. The genome sequence can be obtained from the DNA Data Bank in Japan (Accession number: DRA005797; Bioproject: PRJDB5781).

### Transcriptome analysis

To search for *doublesex* (*dsx*) and *vitellogenin* (*vtg*) homologs, we performed RNA-seq analysis. Adults of 15 ♀♀ and 15 ♂♂ of *T. domestica* were sampled 1440 minutes after a molt in December, 2019. The fat bodies of the individuals were removed using tweezers in a phosphated buffered saline (PBS; pH=7.2). Three adults were used per sample. Total RNA was extracted from 10 samples (5♀♀, 5♂♂) using RNeasy Micro kits (QIAGEN K.K., Tokyo, Japan) following the manufacturer’s instructions. The concentration of purified RNA was measured using a Qubit 4 fluorometer (QIAGEN K.K., Tokyo, Japan) with Qubit RNA BR Assay kits (QIAGEN K.K., Tokyo, Japan). Paired-end libraries were constructed from 100 ng of the total RNAs using TruSeq RNA Library Prep kits v2 (Illumina K.K., Tokyo, Japan) following the manufacturer’s instructions. The libraries were run on a sequence (Hiseq, Illumina, Tokyo, Japan). The library preparation and sequencing were performed by Genewiz Strand-Specific RNA-seq service. We mapped the reads obtained to the assembled genome using the HISAT2 program (Kim et al. 2019) with a default option and counted the mapped reads using the STRINGTie program (Pertea, 2015) with default parameter settings. Differential expression gene analysis was performed based on the count matrix using the “edgeR” package (Robinson et al. 2010) in R-v4.0.3 (*R Core Team, 2020*). Information about the samples can be obtained from the National Center for Biotechnology Information (NCBI) BioSample database (Accession number: SAMN18175012–SAMN18175021).

### Molecular phylogenetic analysis

Dsx is a member of the Doublesex and Mab-3 Related transcriptional factors (DMRT) family, and has a DNA binding domain, Doublesex and Mab-3 (DM) domain. Pancrustacea generally has four DMRT family genes, Dsx, Dmrt11, Dmrt93B, and Dmrt99B (Mawaribuchi et al. 2019). Phylogenetic analysis of Dsx homologs was performed using the amino acid sequences of the DM domain. We used the Dsx sequences of *D. melanogaster* as a query and obtained 97 metazoan DMRT family proteins from the NCBI and the i5k databases (https://i5k.nal.usda.gov/) and our genome data of *T. domestica* by the BLAST analysis (listed in supplementary table 1). We then aligned the sequences using MAFFT version 7 (Katoh et al. 2013) with the -linsi option (to use an accuracy option, L-INS-i) and manually extracted the DM domain, which consisted of 61 amino acids (supplementary fig. 8). The result of the multiple sequence alignment can be obtained from supplementary sequence file 1. Molecular phylogenetic analysis of the aligned sequences was performed using a maximum likelihood method after selecting a substitution model (JTT matrix-based model) with MEGA X (Kumar et al. 2018). Bootstrap values were calculated after 1000 replications.

### Full-length cDNA and exon-intron structures

To elucidate the exon-intron structures of Dsx and Dsx-like, we determined the full-length cDNA sequences using a Rapid Amplification of cDNA Ends (RACE) method and performed a BLAST analysis for our genome database of *T. domestica*. We extracted total RNA from eggs, whole bodies, fat body, and gonads of nymphs and adult females and males of *T. domestica* using TRI Reagent (Molecular Research Center Inc., Ohio, USA) following the manufacturer’s instructions. The total RNAs were treated with RNase-Free DNase I (New England BioLabs Japan Inc., Tokyo, Japan) to exclude remaining genomic DNA and purified by phenol/chloroform extraction and ethanol precipitation. For 50-RACE analysis, mRNAs were purified from 75 µg of the total RNAs using Dynabeads mRNA Purification kit (Thermo Fisher Scientific K.K., Tokyo, Japan) following the manufacturer’s instruction. We then ligated an RNA oligo at the 5’-end of the mRNA using GeneRacer Advanced RACE kits (Thermo Fisher Scientific K.K., Tokyo, Japan). For 30-RACE analysis, we ligated an RNA oligo of the SMART RACE cDNA Amplification Kit (Takara Bio Inc., Shiga, Japan) at 3-end of the total RNA during reverse transcription. First stranded (fs-) cDNA was generated from the RNAs using SuperScript III Reverse Transcriptase (Thermo Fisher Scientific K.K., Tokyo, Japan). We used primers specific to the RNA oligos and performed RACE analysis by nested RT-PCR using Q5 High-Fidelity DNA polymerase (New England BioLabs Japan Inc., Tokyo, Japan). The primers specific to *dsx* and *dsx-like* were made from sequences of the relevant genomic regions and are listed in supplementary table 9. The amplicons were separated using the agarose gel-electrophoresis and cloned using TOPO TA Cloning Kit for Sequencing (Thermo Fisher Scientific K.K., Tokyo, Japan) following the manufacture’s protocol. We used a DH5α *Escherichia coli* strain (TOYOBO CO., LTD., Osaka, Japan) as the host cell. Plasmids were extracted using the alkaline lysis and purified by phenol-chloroform and ethanol precipitation. The nucleotide sequences of the cloned amplicons were determined from the purified plasmids by the Sanger Sequencing service of FASMAC Co. Ltd. (Kanagawa, Japan). We then searched the genomic region of the full-length cDNA sequences of *dsx* and *dsx-like* via local blastn analysis.

### Reverse transcription-quantitative PCR (RT-qPCR)

To quantitative mRNA expression levels, we performed RT-qPCR analysis. For investigating the sex-specific expression profile of *dsx* and *dsx-like*, we used the fat body of adults of *T. domestica* since the sexes can be distinguishable by the external morphology at this stage. Fat bodies also exhibit sex-specific physiological functions in adults. Thirteenth instar individuals and adults after molting were sampled for investigating roles of the genes in the sexually dimorphic morphology and the *vitellogenin* expression, respectively. The sample sizes are reported in the figure legends and supplementary table 2. We dissected the individuals in PBS and collected their fat body in 2 ml tubes containing TRI Reagent (Molecular Research Center Inc., Ohio, USA). The fat bodies then were disrupted using a TissueLyser LT small beads mill (QIAGEN K.K., Tokyo, Japan). These disrupted samples were preserved at −80°C until used. Total RNA was extracted from the samples according to the manufacture’s protocol for the TRI Reagent. Extracted RNA was treated with 2% RNase-free DNase I (New England BioLabs Japan Inc., Tokyo, Japan) at 37°C for 40 minutes and purified by phenol/chloroform extraction and ethanol precipitation.

We measured the concentration of the total RNA using a spectrophotometer (DS-11+, Denovix Inc., Wilmington, USA). fs-cDNA was synthesized from 350 ng of the total RNA using SuperScript III Reverse Transcriptase (Thermo Fisher Scientific K.K., Tokyo, Japan). We diluted the fs-cDNA to 1:2 with MilliQ water and preserved it at −30°C until it was used in RT-qPCR assay. The RT-qPCR assays were performed using a LightCycler 96 instrument (Roche, Basel, Switzerland) according to the manufacture’s protocol with the THUNDERBIRD SYBR qPCR Mix (TOYOBO Co. Ltd., Osaka, Japan). The reaction volume was 10 µl. We used 1 µl of the fs-cDNA as templates. The preparation of the RT-qPCR solution proceeded on ice. The protocol of the RT-qPCR was as follows: preincubation at 95°C for 600 seconds and 45 cycles of three-step reactions, such as denaturation at 95°C for 15 seconds, annealing at 60°C for 15 seconds and extension at 72°C for 45 seconds. We used *ribosomal protein 49* (*rp49*) as a reference gene, as described by Ohde et al. (2011). We designed primer sets of the target genes by the Primer3Web version 4.1.0 (Untergasser et al. 2012) following the manufacture’s recommended condition of the THUNDERBIRD SYBR qPCR Mix. We confirmed the primers’ specificity using melting curves ranging from 65°C to 95°C. We selected primer sets exhibiting a single peak. The primers are listed in supplementary table 9. Each RT-qPCR was technically replicated three times.

Some samples were excluded before analyzing the data when the Ct value of any genes was not detected in one or more replicates or when the Ct value of the reference gene deviated from that of other samples. In these removed data, a technical error was suspected. We calculated the expression level of target genes by the 2^−ΔΔCt^ method (Livak and Schmittgen 2001) and performed the Brunner–Munzel (BM) test for ΔCt value. The BM test was carried out using R-v4.0.3. with the *brunnermuzel.test* function of the “brunnermuzel” package (https://cran.r-project.org/web/packages/brunnermunzel/index.html). Holm’s method was used for multiple comparison analyses between the control and treatments. The data are listed in supplementary table 2. In the *dsx* expression of the RNAi male, we performed the Smirnov-Grubbs (SG) test for ΔCt value using the *grubbs.test* function of the “outliers” package in R (https://cran.r-project.org/web/packages/outliers/index.html) (supplementary table 3). An outlier was detected in the *dsx* RNAi male. We repeatedly performed the SG test using the data excluding the outlier. No further outliers were detected. Lastly, we re-analyzed the data, excluding the outlier, using the BM test (supplementary table 2).

### RNAi analysis

The RNAi assay can be used to examine the roles of genes during postembryonic development in *T. domestica* (Ohde et al. 2011). The sexual differentiation of insects is generally assumed to be a cell-autonomous mechanism that is independent of systemic hormonal-control (Verhulst and van de Zande 2015) as discussed in De Loof and Huybrechts (1998) and Bear and Monteiro (2013) and progresses during postembryonic development. Therefore, nymphal RNAi is the most effective tool to investigate the roles of genes on sexual trait formation during postembryonic development. To reduce the risk of off-target effects, the dsRNA was designed to avoid the region of the DM domain. We also confirmed that the dsRNA had no contiguous matches of more than 20 bases with other genes on the genome by BLAST (blastn option). To produce templates for the dsRNA, we cloned the regions of *dsx* and *dsx-like* from the fs-cDNA using the same method as the RACE analysis. We amplified the template DNAs from purified plasmids with PCR using Q5 High-Fidelity DNA Polymerase and purified the amplified DNA with the phenol/chloroform extraction and the ethanol precipitation. dsRNA was synthesized from the purified DNA using Ampliscribe T7-Flash Transcription kits (Epicentre Technologies, Co., Wisconsin, USA). We designed the PCR primers using the Primer3Web version 4.1.0 (Untergasser et al. 2012). The PCR primers are listed in supplementary table 9. In nymphal RNAi analysis, we injected the dsRNAs repeatedly into the abdomen of the nymphs of *T. domestica* with each molt from the fourth or fifth instar to thirteenth instar to sustain the RNAi effect during postembryonic development. The initial stage was the same within a single experiment. This repeated RNAi treatment was effective in some insects such as *Blattella germanica* (Wexler et al. 2019). We sampled the individuals one, three, and five days after molting, using phenotypic observations, analysis of *dsx* knockdown effects, and the oocyte number.

To determine the sex of individuals, we initially observed the gonads: testis and ovary. In our RNAi analysis, the gonads completely formed and there was no difference between the control and *dsx* RNAi individuals in external morphology (fig. 2C). Therefore, individuals with testis were males and those with ovaries were females. *T. domestica* molts throughout its life, even after sexual maturation, and produces *vtg* during each adult instar (Rousset and Bitsch 1993). To analyze the *vtg* mRNA levels, we also injected the dsRNAs of *dsx* and *dsx-like* repeatedly into the females and males every three days from 12 hours after molting. We sampled the females and males at 720±20 minutes after subsequently molts.

### Phenotype observation

We dissected thirteenth instar individuals in PBS using tweezers and removed the thoraxes, reproductive systems, and external genital organs. We took images using the digital microscope system (VHX-5000, KEYENCE, Tokyo, Japan). The thoraxes and external genital organs were fixed with FAA fixative (formaldehyde: ethanol: acetic acid = 15:5:1) at 25°C overnight and then preserved in 90% ethanol. We used the length of the prothorax as an indicator of body size. To measure the prothoracic width, the prothoracic notum was removed from the fixed thorax after treatment with 10% NaOH solution at 60°C for 30 minutes to dissolve the soft tissues. The notum was mounted in Lemosol on a microscope slide. The prepared specimens were imaged using a KEYENCE VHX-5000. With the microscope at 50×, the length of the notum was measured. The ovipositor length was also measured using the microscope at 20× and 50×. To count the sperm number, sperm was collected from seminal vesicles and diluted with 5 ml MilliQ water. 50 µl of the diluted sperm was spotted on a microscope slide and dried overnight. We technically replicated the measurement three times for ovipositor length and six times in sperm number and calculated these means. Measurement was performed by blinding the treatment. We counted the number of oocytes in ovarioles using an optical microscope at 50× (Olympus, Tokyo, Japan). A generalized linear model (GLM) was used to analyze differences in ovipositor length (length data) and sperm and oocyte number (count data) among RNAi treatments. The body size, target genes, and interactions between the target genes were used as explanatory variables. The length was assumed to follow a Gaussian distribution, and the count data to have a negative binomial distribution. We used R-v4.0.3 in these analyses and the *glm* and the *glm.nb* (MASS package) functions for the length and count data, respectively. To analyze the contribution of the explanatory variables, a likelihood ratio test for the result of GLM was performed using the *Anova* function of the car package. The statistical results are listed in supplementary table 4 (female) and 5 (male).

### Scanning Electron Microscopy (SEM)

The NanoSuit method (Takaku et al. 2013) was used for the SEM analysis. Male penises and female ovipositors preserved in 90% ethanol were washed with distilled water and immersed in 1% Tween20 at 25°C for 10 minutes. The samples were mounted on stubs and imaged using a low-vacuum SEM (DX-500; KEYENCE, Tokyo, Japan).

### Histology

The gonads of RNAi individuals were fixed with Bouin’s fixative (saturated picric acid: formaldehyde: glacial acetic acid = 15:5:1) at 25°C overnight and washed with 90% ethanol plus Lithium Carbonate (Li_2_CO_3_). The ovipositors of RNAi individuals were fixed with FAA fixative at 25°C overnight and then were transferred into 90% ethanol. The samples were dehydrated and cleared with an ethanol-butanol series. The cleared samples were immersed and embedded in paraffin at 60°C. The paraffin blocks were polymerized at 4°C and cut into 5 µm thick sections using a microtome (RM2155: Leica, Wetzlar, Germany). The sections were mounted on microscope slides coated with egg white-glycerin and stained using Delafield’s Hematoxylin and Eosin staining. After staining with the hematoxylin, the slides were washed with 1% hydrochloric acid-ethanol for 40 seconds. The stained slides were enclosed with Canada balsam. We observed the slides on an optical microscope (Olympus, Tokyo, Japan) and took photos using a digital single-lens reflex camera (Nikon, Tokyo, Japan).

### Ancestral Sequence Reconstruction

To infer the sequence evolution of the *dsx*, we conducted an ancestral sequence reconstruction (ASR) of the C-terminal sequences of the *dsx* female-type homologous sequence. First, we searched homologous sequences to *dsx* female-type from NCBI protein/transcript shotgun assembly databases and previous studies. The searches in the NCBI databases were performed by BLAST search. We closely examined the alignment results of the BLAST and selected sequences with at least 10 amino acids aligned with the female-specific region of each query sequence. We do not know whether some of these sequences are expressed in females and contribute to female morphogenesis, as these sequences are not necessarily to have investigated expression and function in the species. We decided that it was not problem to use these sequences since we focused on the evolution of sequences homologous to *dsx* female-type in each insect taxa. In Diptera, we set *dsx* female-type of *D. melanogaster* (Accession #: NP_001287220) as a query and obtained 9 sequences. In Lepidoptera, we used *dsx* female-type of *B. mori* (NP_001036871) as a query and get 10 sequences. In Coleoptera, *dsx* female-type of *Tribolium castaneum* (AFQ62106) was set in a query and then 10 sequences were obtained. We used *dsx* female-type of *Ap. mellifera* (NP_001128407) and *At. rosae* (XP_012262256) as queries to search hymenopteran sequences. We also searched some hymenopteran sequences from the NCBI databases based on a previous study (Baral et al. 2019). 10 hymenopteran sequences were obtained. In Psocodea and Hymenoptera, we searched the databases to set the sequences of *Pediculus humanus* (QGB21102) and *Rhodonius prolixus* (QGB21099) as queries. Wexler et al. (2019) showed that *dsx* of *Pediculus humanus* (Psocodea) has isoforms without sex-specificity. In this study, based on the blast search and exon structure, we regarded that the PhDsx1 in Wexler et al. (2019) is homologous to the *dsx* female-type. The sequences of *Ni. lugens* (AWJ25056) and *Bl. germanica* (QGB21105 and QGB21106) were obtained from the database based on previous studies (Zhuo et al. 2018; Wexler et al. 2019). We selected two sequences from *Bl. germanica*, as this species has two female-specific *dsx* isoforms (Wexler et al. 2019). The sequences of *Cryptocercus punctulatus* and *I. senegalensis* were obtained from previous studies (Miyazaki et al. 2021; Takahashi et al. 2021). In *T. domestica*, the sequence identified in this study was used. The sequence names are listed in supplementary table 7. We then manually extracted the OD domain and performed multiple sequence alignments (MSA) using the MAFFT version 7 (Katoh et al. 2013) with the -linsi option (to use an accuracy option, L-INS-i) (supplementary sequence file 2). We reconstructed ancestral sequences (AS) from the MSA using MEGA X software. The maximum-likelihood method was applied to the ASR. The JTT + G model was chosen as a substitution model by AIC-based model selection. The guide tree was reconstructed based on previously reported phylogenetic relationships (Wiegmann et al. 2011; Misof et al. 2014; Li et al. 2017; Peters et al. 2017; Zhang et al. 2018; Kawahara et al. 2019; McKenna et al. 2019; Gustafson et al. 2020) (supplementary fig. 9). We selected the most probable sequences for the following analyses. The results of ASR can be seen in supplementary table 8. The probabilities of sites of AS that we focused on are listed in supplementary table 10. In Aparaglossata (Node 77) and Holometabola (Node 87) AS, almost all probabilities of sites were more than 0.9. The except sites were sites 83 and 98 in Node 77 and sites 77–79 and 83 in Node 87. These sites other than sites 77 had probabilities > 0.5. Thus, we concluded that the AS in Aparaglossata and Holometabola, which we considered the most critical, was reconstructed with sufficient reliability. Any residues had the probabilities = 0 in the Aparaglossata-specific region of Holometabola AS. In contrast, in non-holometabolan insects, since our taxon sampling is limited to several species (Eumetabola in Node 92, Neoptera in Node95, Pterygota in Node 96), the probabilities of some sites are lower than 0.5. These low probable sites are not necessarily confident. To conclude with reliability, it is no doubt that analyses based on a larger number of species will be essential. However, all sites of the Aparaglossata-specific region in these AS were gaps with the probabilities > 0.9. The result of the sites of the Aparaglossata-specific region seems to be relatively reliable in our analysis. Thus, our conclusion that the Aparaglossata-specific region occurred in the common ancestor of Aparaglossata would be confident. To compare the sequences, we then performed MSA of the most probable reconstructed ancestral sequences and the sequence of *D. melanogaster* using MAFFT version 7 (fig. 5A).

### Protein Structure Prediction

To infer the evolution of protein structures of *dsx*, we conducted the protein structure prediction. The ancestral sequences reconstructed by the above section were used for the protein structure prediction. The sequences were obtained from supplementary sequence file 3. The protein structure prediction was performed using the Alphafold2-based algorism (ColabFold: Mirdita et al. 2021) with the default option. The accuracy of predictions was evaluated based on the predicted Local distance difference test (plDDT) score that was automatically calculated on the ColabFold. We selected a model with the highest average plDDT score in each prediction. The average plDDT scores were 81.824 (Aparaglossata), 89.165 (Holometabola), 87.376 (Eumetabola), 90.721 (Neoptera), and 90.720 (Pterygota). The plDDT scores were more than 70 in the helical structure predicted as the α-helix loop of the female-specific *dsx* region. Generally, predicted structures of plDDT>70 are regarded to be a confident prediction (cf., Tunyasuvunakool et al. 2021).

Therefore, we assessed the α-helix loop of the female-specific region of *dsx* as the confidently predicted structure. The graph of the plDDT score of each model is shown in supplementary fig. 10. The 3D models of predicted structures were visualized with the PyMOL Molecular Graphics System, Version 2.0 (Schrödinger, LLC.). On the viewer, we colored the female-specific region and the Aparaglossata-specific region with red color and the green color, respectively.

### Data availability

The draft genome data was deposited in the DNA Data Bank of Japan (Accession number: DRA005797; Bioproject: PRJDB5781). The raw read data of the transcriptome was in the NCBI Sequence Read Archive (Accession numbers: SRR13870115–SRR13870124; Bioproject: PRJNA707122). The sequences of *dsx* male-type, *dsx* female-type, and *dsx-like* are also in GenBank (Accession numbers: MW711323, MW711324, and MW711325, respectively).

## Acknowledgments

We express gratitude to Dr. Daniel Bopp (University of Zürich) for his comments and encouragement for this manuscript. We would like to thank Dr. Takahiro Ohde (National Institute for Basic Biology; Kyoto University) for his help to extract genomic DNA of *T. domestica* and for a great advice to the discussion. We are also grateful to Dr. Toshiya Ando, Dr. Taro Nakamura, Dr. Shinichi Morita, Dr. Hiroki Sakai, and Dr. Tatsuro Konagaya (National Institute for Basic Biology) for technical advice and discussion on this manuscript. We also express our gratitude to Dr. Satoshi Miyazaki (Tamagawa university) for providing sequences of *dsx* in *Cryptocercus punctulatus*. Computations were performed on the NIG supercomputer at ROIS, National Institute of Genetics and the Data Integration and Analysis Facility, National Institute for Basic Biology. We thank the Model Plant Research Facility, NIBB Bioresource Center for providing the network camera system. This work was supported by the JSPS KAKENHI Grant numbers JP25660265, JP16H02596, and JP16H06279 (PAGS) for TN and the Sasakawa Scientific Research Grant from The Japan Science Society for YC.

## Author Contributions

YC and TN conceived this study. YC performed all experiments, observations, and analyses other than the genome sequence and assembly. AT sequenced the genome. MO and TI performed the de novo genome assembly. YC and TI wrote the manuscript. All authors commented on the manuscript.

## Competing Interest Statement

The authors declare that have no competing interests.

## Notes

### Competing Interest Statement

The authors have declared no competing interest.

