## Supplementary material for "Evolutionary history of sexual differentiation mechanism in insects"

Teruyuki Niimi

**This PDF file includes:**

Supplementary text

Supplementary Figures 1 to 10

Caption of Supplementary Tables 1 to 9

Supplementary references

**Other supplementary materials for this manuscript include the following:**

Supplementary Tables 1 to 9: an EXCEL file

Supplementary Sequence Files: two FASTA format files

**Supplementary Text**

**Detailed results**

The general morphology of the external genital organs and the reproductive systems in Zygentoma is reviewed in Matsuda (1976). The recent interpretation of the penial structure and its evolution are explained in Boudinot (2018). The ovipositor structure and its evolution are detailed in Emeljanov (2014). Here, we described the morphology of the genital organs and the reproductive systems in *Thermobia domestica* with our focus points.

**Reproductive system and germline cell morphology in males**

In males of *T. domestica*, a pair of testes was located on the dorsal side of the abdomen. The testis was consisted of some testicular follicles (fig. 2C, D). Each testicular follicle was connected to the vas deferens via the vas efferens (fig. 2D). The seminal vesicle lay between the vas deferens and the ejaculatory duct. A pair of the ejaculatory ducts was associated with each other in the front of the gonopore in the penis (fig. 2D). The testicular follicles were a bean-like shape and the seminal vesicles were a bean pod-like shape. In the testicular follicle, the spermatogonia was in the antero-most part (fig. 2C). The primary and secondary spermatocytes lay in the middle part. In the posterior part of the testicular follicle, there were some sperm bundles (fig. 2C). The wall of the testicular follicle consisted of a single flattened epithelial layer.

We observed the above features of the reproductive system in *dsx* or *dsx-like* RNAi males (fig. 2C, D). In *dsx* and both genes RNAi males, the seminal vesicles were rounded shape. The vas efferens was filled with the sperm (fig. 2D). In contrast, we could not find differences in the morphology of the testicular follicles or spermatogenesis between the RNAi and control males. The male reproductive system and spermatogenesis showed no visible difference between *dsx* female-type and *dsx-like* RNAi females and the control ones.

**Reproductive system and germline cell morphology in females**

In females of *T. domestica*, part of the ovary was on the dorsal side of the abdomen. Each ovary consists of five ovarioles and was attached to the anterior part of the abdomen via the terminal tuft (fig. 2F). The ovarioles were associated with each other at the lateral oviduct. The lateral oviduct was connected to the common oviduct and subsequently opened at the gonopore in the valvula I. There was no vagina between the gonopore and the oviduct. The spermatheca was located on the branch point of the common oviduct along the midline (fig. 2F). The spermatheca was divided into two parts: anterior and posterior (supplementary fig. 2). The anterior part consisted of a pseudostratified layer of the columnar epithelial cells that were secretory. The posterior part was surrounded by a single layer of epithelial cells. The ovariole was panoistic-type and was composed of two parts: the germarium and the vitellarium fig. 3C). The germarium contained many oogonia and young oocytes. The vitellarium had previtellogenic and vitellogenic oocytes. The oocytes in the vitellarium were surrounded by a single layer of follicle cells. There were pedicel cells in the terminal of the ovariole. The previtellogenic oocyte had a large germinal vesicle and basophilic cytoplasm. The vitellogenic oocyte was elongated along the anterior-posterior axis of the ovariole and had eosinophilic cytoplasm. Many eosinophilic lipid droplets were present in the peripheral region of the vitellogenic oocytes. The follicle cells were flattened and columnar in shape in the previtellogenesis and the vitellogenesis.

We observed the above features of the reproductive system in *dsx* or *dsx-like* RNAi females (fig. 2C, F; supplementary fig. 2). We could not detect visible differences in the female reproductive system or oogenesis between the RNAi females and the controls. This result suggests that the *dsx* and *dsx-like* have no function in the formation of female traits and gametogenesis at the tissue and cellular level.

**General morphology of the external genital organ in *Thermobia domestica***

The description below is based on the observation of the controls, i.e., individuals injected *egfp* dsRNA, and is agreed with previous studies in *T. domestica* (Snodgrass 1957; Matsuda 1976; Emeljanov 2014; Boudinot 2018).

*T. domestica* males have a single penis. This penis is an unpaired appendix on the abdomen segment IX and is not aedeagus (copulatory organ) (Matsuda 1976). The penis was sub-segmented into two parts. There were many setae on the left and the right side of the distal tips (fig. 3C). The surface of the penis had a reticulated pattern (fig. 3C). This simple penial structure was presumably gained at the last common ancestor of Ectognatha (= Archaeognatha + Zygentoma + Pterygota: Insecta s.str.) (*Boudinot, 2018*).

*T. domestica* females has an ovipositor. This ovipositor consists of two pairs of appendices (gonapophysis) and is derived from the retracted vesicles on the abdomen VIII and IX (Matsuda 1976; Emeljanov 2014). This ovipositor is an autapomorphy of Ectognatha (Kristensen 1975; Beutel 2017). The gonapophyses on the abdomen VIII (valvula I) were the ventral part of the ovipositor and a paired structure. The gonapophyses on the abdomen IX (valvula II) were the dorsal side of the ovipositor and were united to form an unpaired structure (supplementary fig. 3B). The distal tip of the valvula II remained a paired structure and possessed dense setae (fig. 3E), which may play a role in sensory reception. Both valvulae were sub-segmented and have some setae (fig. 3E). The valvula I and II were connected through a tongue-and-groove structure (olistheter). The olistheter consisted of an aulax (”groove”) on the valvula I and a rhachis (”tongue”) on the valvula II (supplementary fig. 3). Within the valvulae, the epithelial cells were beneath the cuticular layer. The cuticular layer was thickened and multi-layered in the outer surface of the ovipositor. In contrast, the inner surface (i.e., the side of the egg cavity) of the ovipositor had a thin and single-layered cuticle. Some lumens of the valvulae were extended along the anterior-posterior axis and were hemocoelic cavities.

Effects of knockdown of *dsx* and *dsx-like* on the external genital organ

In *dsx* RNAi males, a tubular organ was formed instead of the penis (fig. 3B, C). This tubular organ consisted of two pairs of appendage-like structures. The inner one is connected to the gonopore and the ejaculatory duct. The outer one had a lot of setae on its tip (fig. 3C). Thus, the inner pair was similar to the valvula I of the female ovipositor and the outer one was similar to the valvula II. We could detect sub-segmentation in both structures (fig. 3C). These features indicated that the tubular organ in the *dsx* RNAi males was parallel to the female ovipositor. The same phenotype was found in the *dsx* and *dsx-like* double RNAi males. In contrast, the *dsx-like* males possessed a penis the same as that of the control insects.

In females that were treated with RNAi for *dsx*, *dsx-like*, and both genes, the external genital organ was the same as the ovipositor of the control females that described in the above section (fig. 3E, F). This genital organ of RNAi females consisted of two pairs of sub-segmented appendage-like structures and possessed dense setae on the tip of the inner pair. The outer pair was connected to the gonopore and the common oviduct. Thus, in the view of histology, the location, and the relation to other elements, the external organ of the RNAi females was not different from the ovipositor of the control ones.

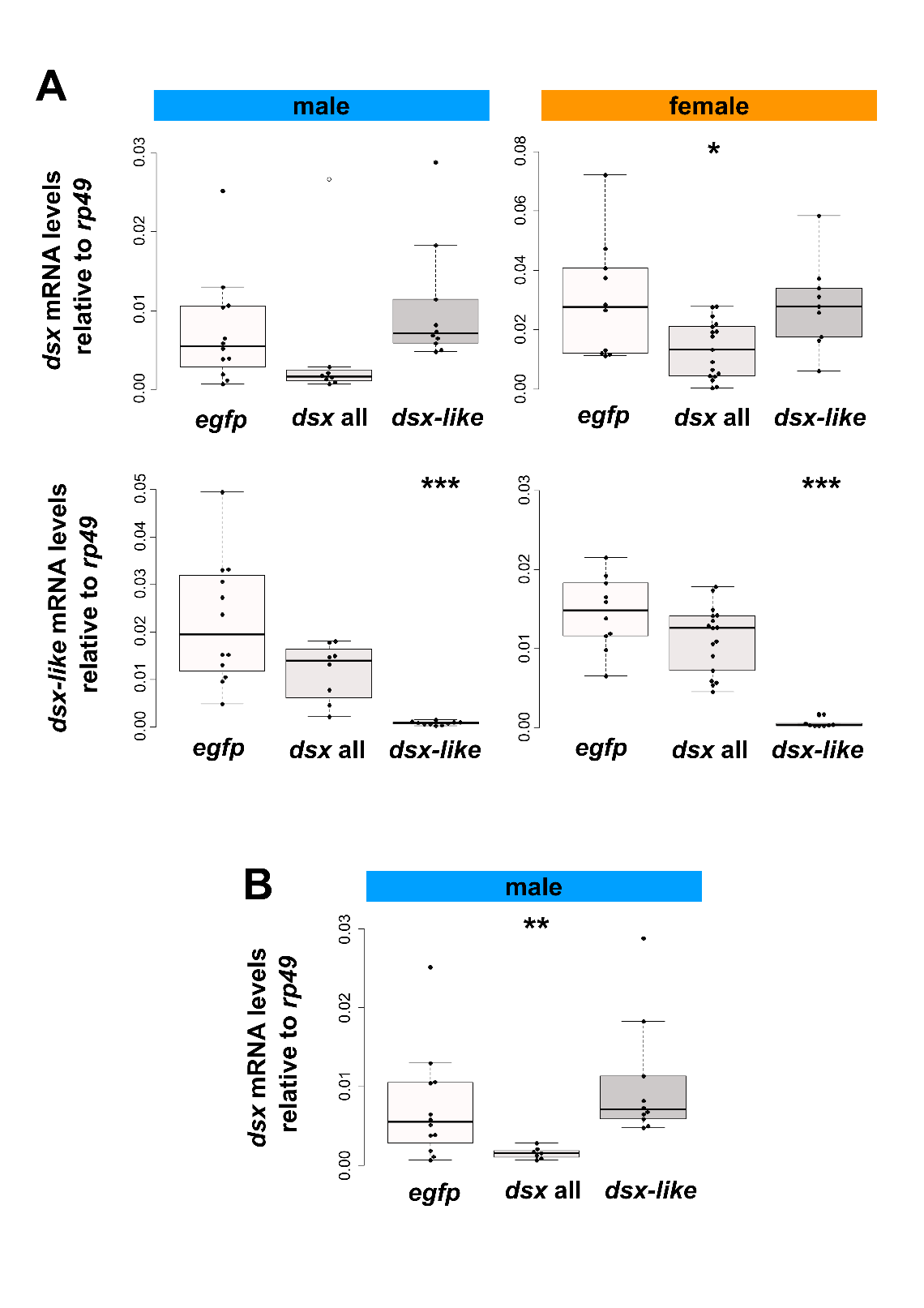
**Supplementary FIG. 1.** Expression of *dsx* and *dsx-like* mRNA in nymphal RNAi individuals. (*A*) Expression level of target genes in RNAi individuals. The mRNA levels of *dsx* and *dsx-like* were analyzed by RT-qPCR assay and are the relative value to the expression of the reference gene, *ribosomal protein 49* (*rp49*). The upper graphs are the expression of *dsx* and the lower ones are that of *dsx-like*. The left column is the result in males and the right one is that in females. White plot suggests the outlier. (*B***)** the expression level of *dsx* mRNA in the nymphal RNAi males after excluding an outlier. To test the outlier, the Smirnov–Grubbs’ test was performed. The result of the Smirnov–Grubbs’ test is shown in supplementary table S3. The *egfp*, *dsx* all and *dsx-lik*e indicates the *egfp* dsRNA injected group (control), *dsx* sex-common region dsRNA injected group and *dsx-like* dsRNA injected group, respectively. Results of the Brunner–Munzel test are indicated by asterisks: **P* < 0.05; ***P* < 0.01; ****P* < 0.001 and is also described in supplementary table S2. *P* ≥ 0.05 is not shown. Each plot indicates the value of each individual. Total *N* = 30 and 36 in males and females.

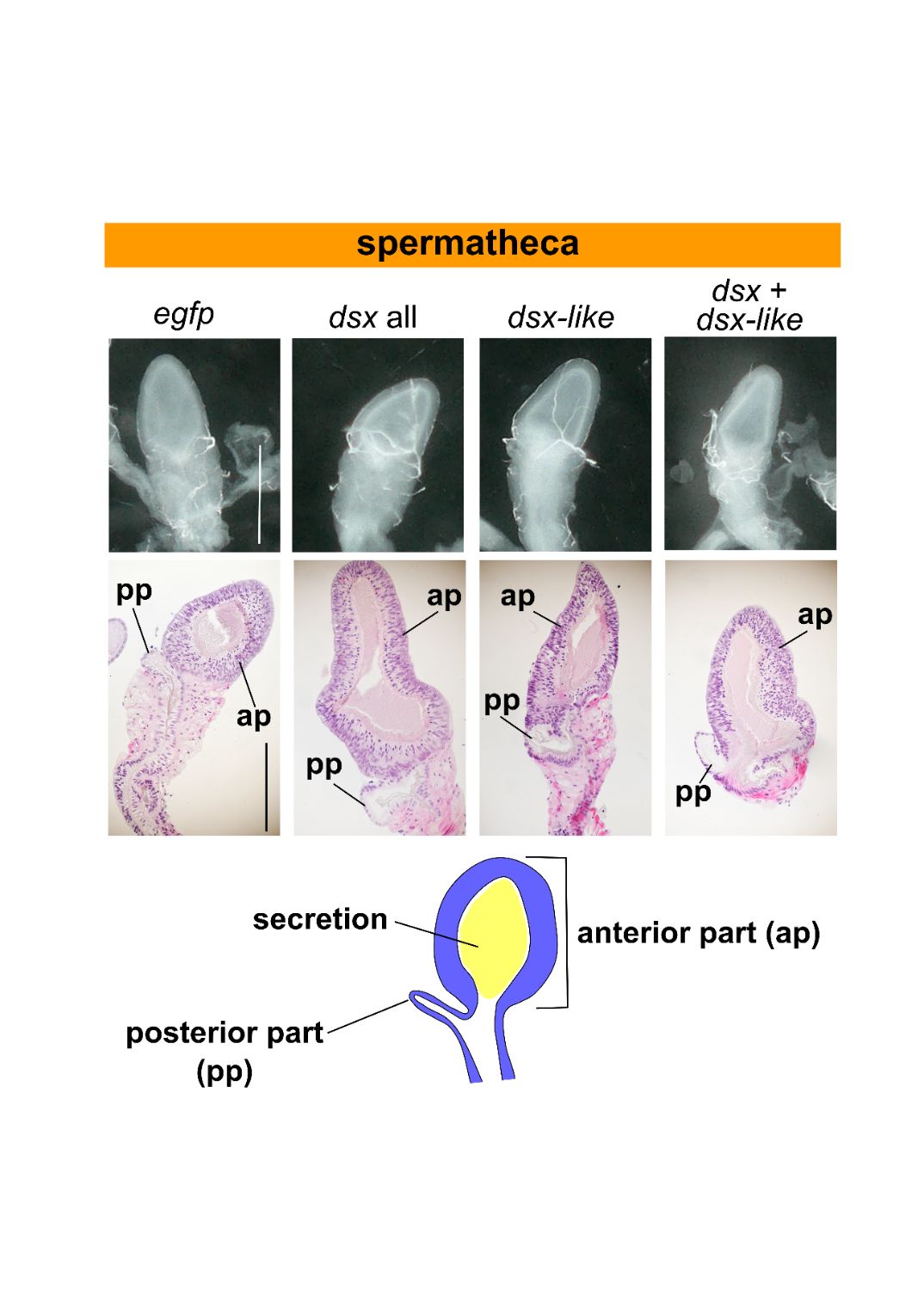
**Supplementary FIG. 2.** Morphology of spermatheca in nymphal RNAi females. The upper photos show the light microscopic images of the spermatheca. The middle ones are paraffin sections of the spermatheca. Hematoxylin-Eosin staining. The lower one is the schematic image of the spermatheca of *T. domestica*. The *egfp*, *dsx* all, *dsx-lik*e and *dsx*+*dsx-like* indicates the *egfp* dsRNA injected group (control), *dsx* sex-common region dsRNA injected group, *dsx-like* dsRNA injected group, and both *dsx* sex-common region and *dsx-like* dsRNAs injected group, respectively. The spermatheca is divided into two parts: anterior and posterior part and have secretion within its lumen. Scales: 500 µm. ap, anterior part of spermatheca; pp, posterior part of spermatheca. The detailed description can be seen in Supplementary Material.

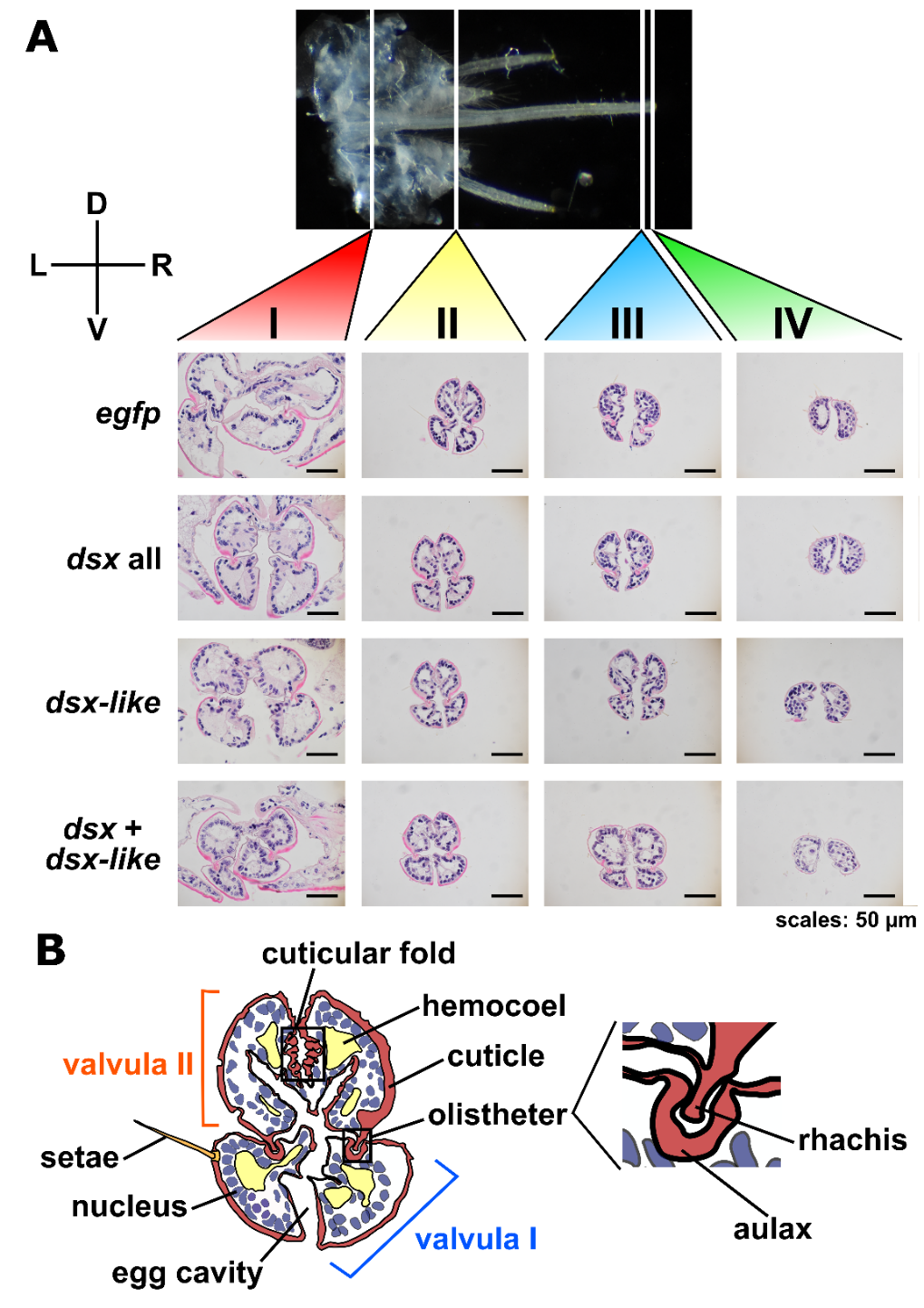
**Supplementary FIG. 3.** Morphology of ovipositor in nymphal RNAi individuals. (A) Cross-section of the ovipositor. The photos show the morphology of the ovipositor in four parts: I (proximal part), II (middle part), III (distal part), and IV (most-distal part). The *egfp*, *dsx* all, *dsx-lik*e and *dsx*+*dsx-like* indicates the *egfp* dsRNA injected group (control), *dsx* sex-common region dsRNA injected group, *dsx-like* dsRNA injected group, and both *dsx* sex-common region and *dsx-like* dsRNAs injected group, respectively. D, dorsal; L, left; R, right; V, ventral. Paraffin. Hematoxylin-Eosin staining. Scales: 50 µm. (B) Schematic figure of the ovipositor morphology. This figure is based on the cross-section of the part II in the control female. The part of ovipositor is constituted of two regions: valvula I and II. These regions are coordinated at the olistheter. The dorsal side of valvula II has folded cuticle. The detailed description can be seen in Supplementary Material.

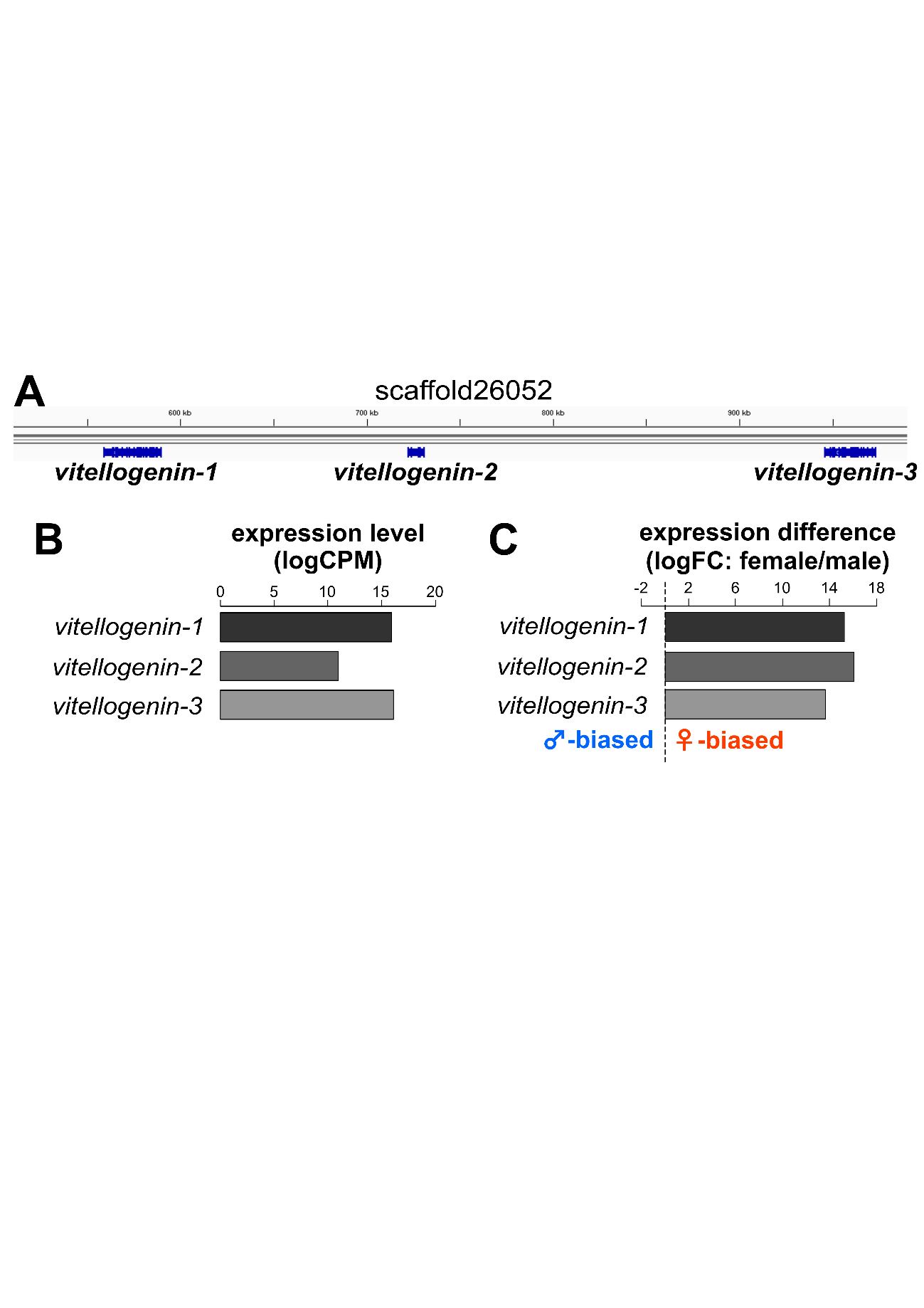
**supplementary FIG. 4.** Expression of *vitellogenin* homologs in *T. domestica*. (*A*) Genome mapping of *vitellogenin* mRNA sequences. The picture is a screenshot of the integrative genome viewer (IGV). Vitellogenin genes are located on scaffold26052 of the assembled genome. *T. domestica* has three tandem-repeated *vitellogenin* homologs: *vitellogenin-1*, *vitellogenin-2*, and *vitellogenin-3*. (*B*) Expression level of *vitellogenin* homologs. The expression levels were calculated by the logCPM of transcriptome data in the fat body of males and females. All *vitellogenin* homologs show the high expression level in the fat body. (*C*) Difference in expression of *vitellogenin* homologs between sexes. The differential expression analysis was performed using the edgeR program. The expression difference is shown by the logFC value. When the values are more than 0, genes are expressed higher in females than males. Thus, all *vitellogenin* homologs shown here are expressed much higher in females than in males. Each value in (*B*) and (*C*) can be seen in supplementary table S6.

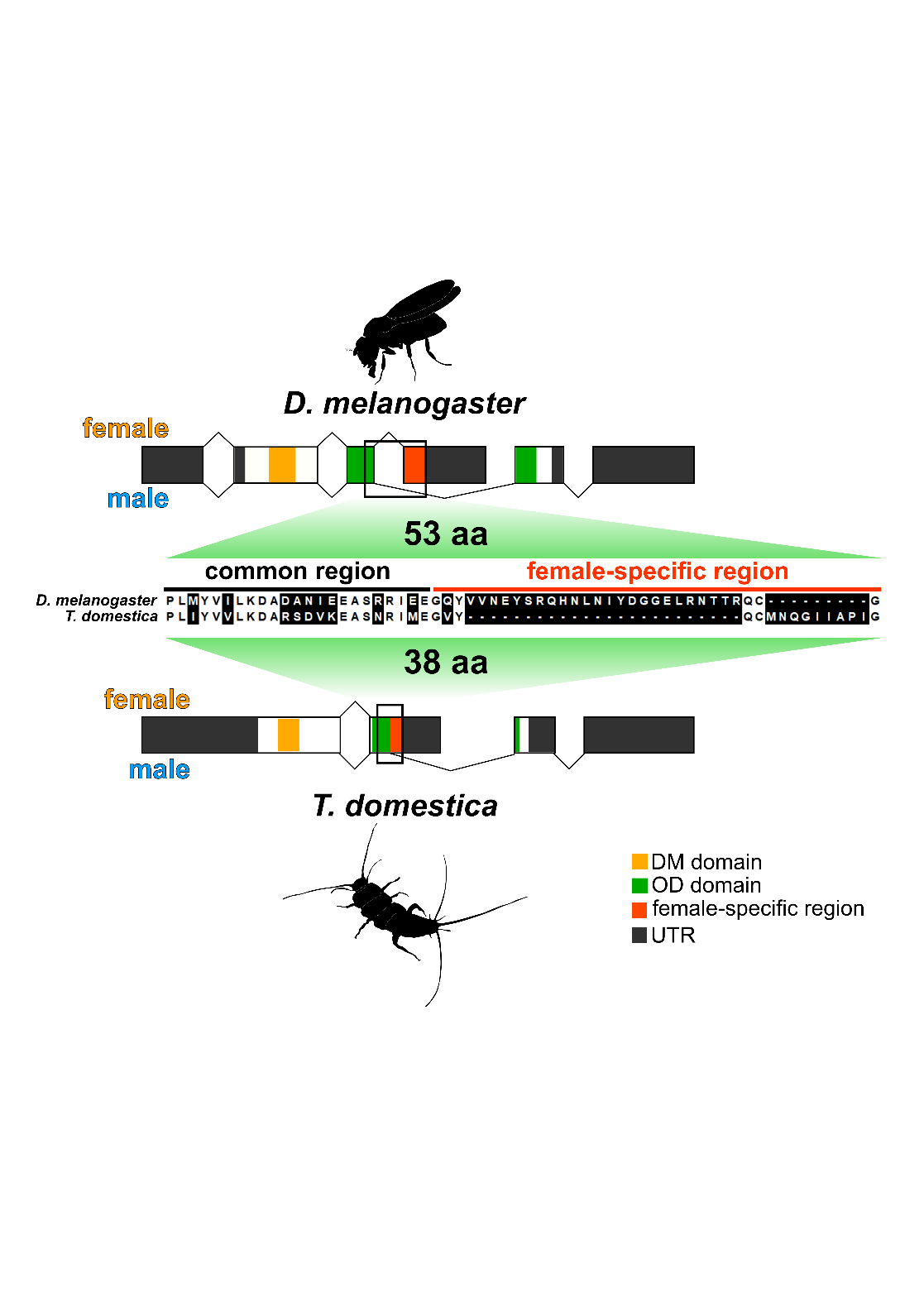
**Supplementary FIG. 5.** Comparison of C-terminal sequences of *dsx* female-type between *Drosophila melanogaster* and *Thermobia domestica*. The upper schematic figure shows the gene structure of *dsx* in *D. melanogaster*. The lower schematic figure indicates the gene structure of *dsx* in *T. domestica*. The female-specific region is shown by the orange color. The middle image is the result of the multiple sequence alignment (MSA) of C-terminal region between two species. The MSA was performed using the MAFFT software. The white background indicates the matched residues between the species in the MSA. The female-specific region is much shorter in *T. domestica* (38 aa) than in *D. melanogaster* (53 aa).

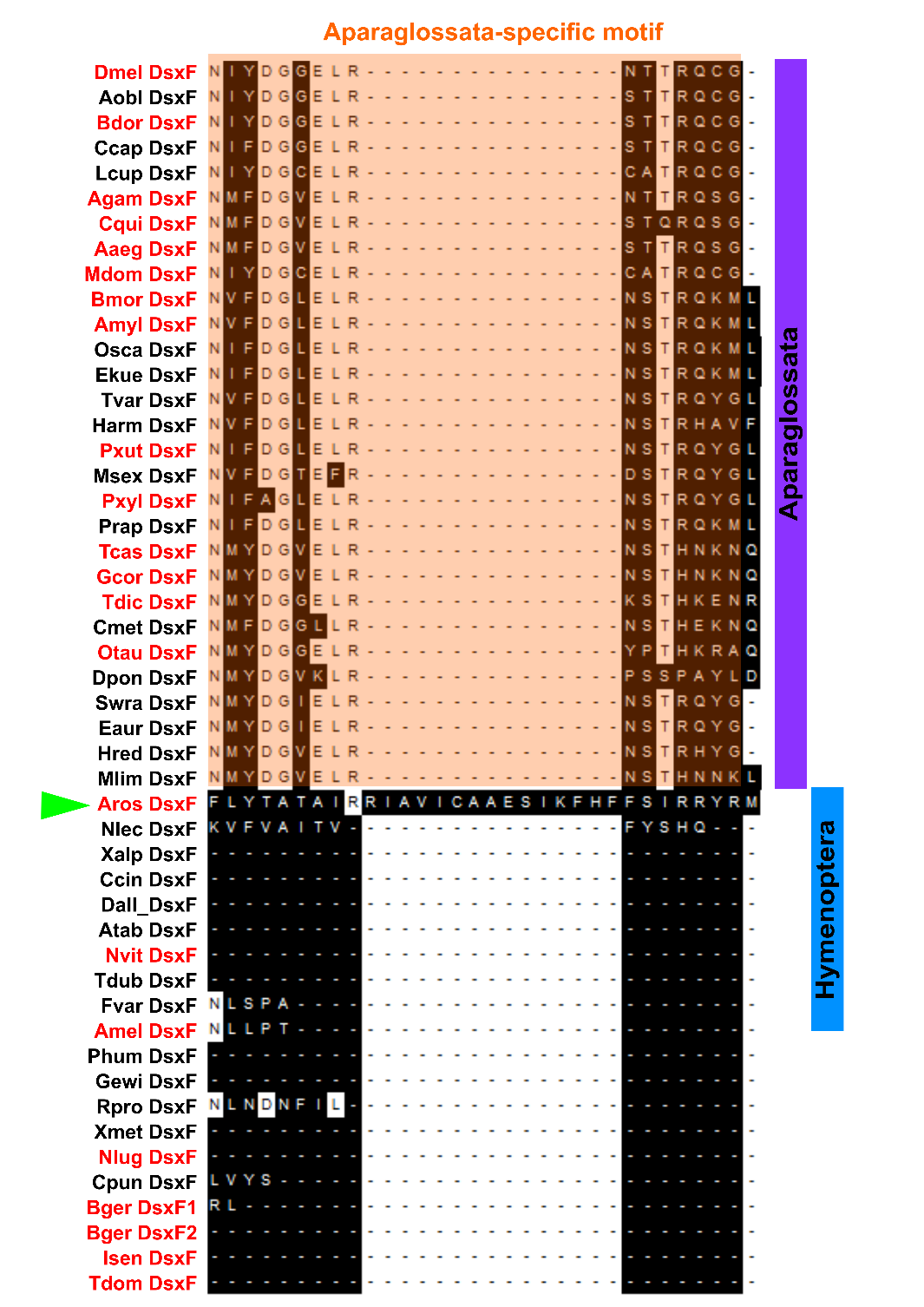
**Supplementary FIG. 6.** Multiple sequence alignments of the Aparaglossata-specific motif. The multiple sequence alignment of the C-terminal sequences was performed by the MAFFT software. This image shows the region around the Aparaglossata-specific motif indicated by the orange color. The species in the sequence name can be obtained from the supplementary table 7. The names indicated by the red color show species in which the functional analysis of *dsx* has been performed. The green arrowhead exhibits the sequence of *Athalia rosae* that has the amino acid sequences corresponding to the motif. The full result of the multiple sequence alignment can be obtained from the supplementary sequence file 2 (FASTA format).

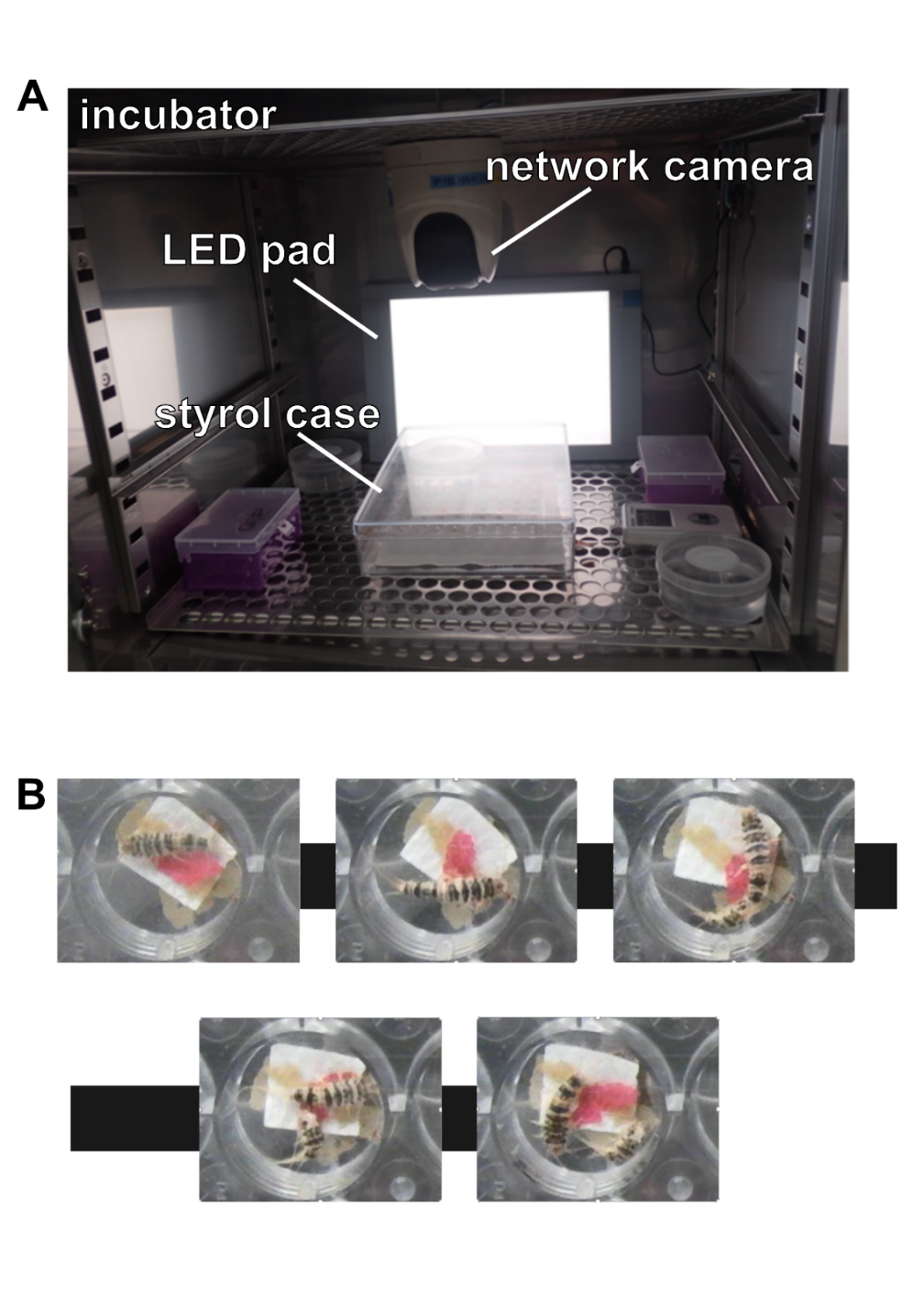
**Supplementary FIG. 7.** Time-lapse imaging system. (*A*) A photo of the time-lapse imaging system used to observe the molt of *T. domestica*. The network camera is located on the floor of the incubator. The insects are put on 24-well plates in the styrol case. The LED pad is used for lighting up the inside of the incubator. The temperature within the incubator is kept at 37°C. (*B*) The time-lapse images during the molt. We set the interval of taking a photo every 5 minutes. The photos are ordered along with the time course from upper-right to lower-left. The firebrat proceeded its molting for 10 minutes.

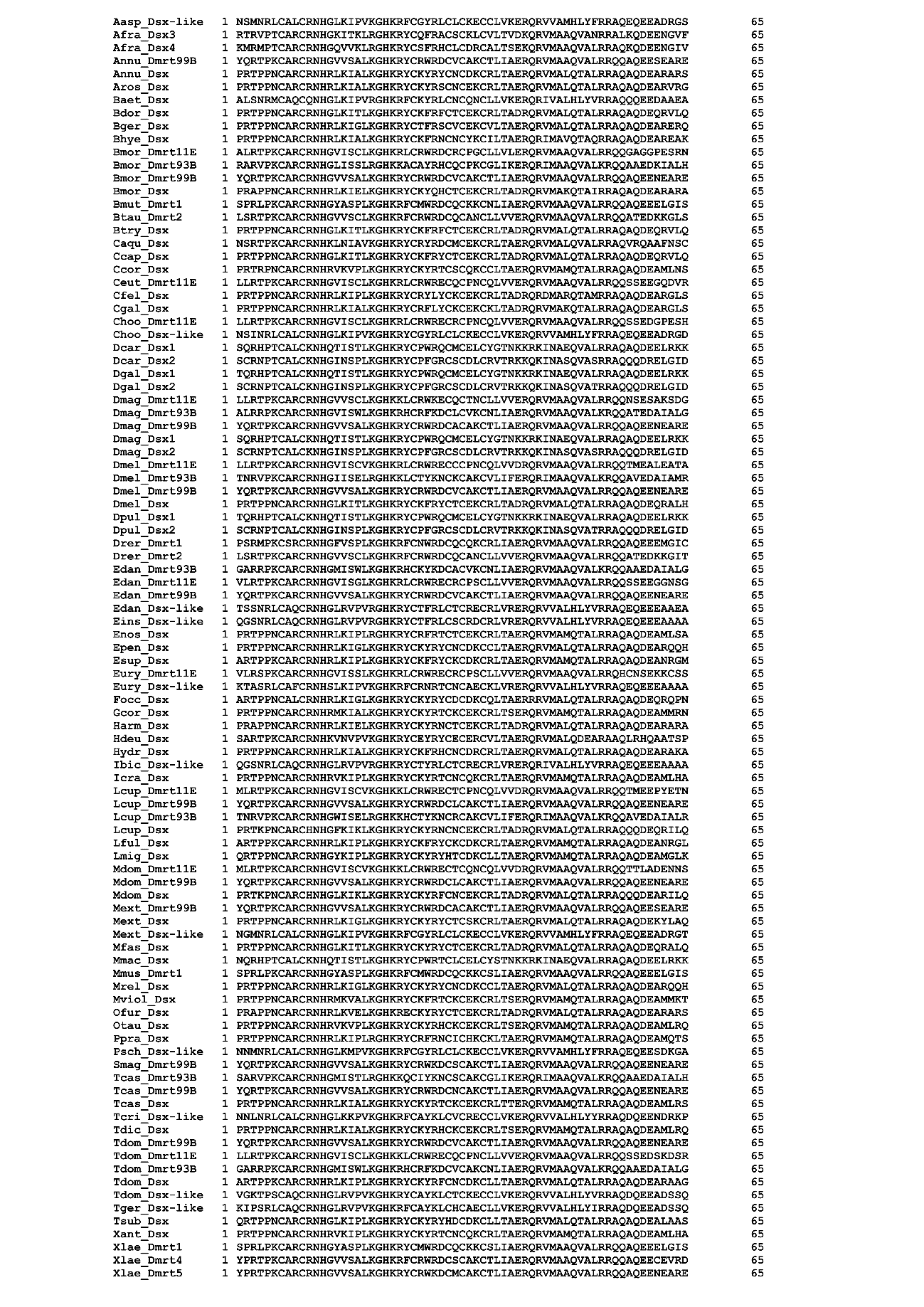
**Supplementary FIG. 8.** Multiple sequence alignment of DM domain of DMRT family proteins for molecular phylogenetic analysis. The 65 amino acids of 97 DMRT proteins were used for the molecular phylogenetic analysis. The sequence file can be obtained from the supplementary sequence file 1 (FASTA format).

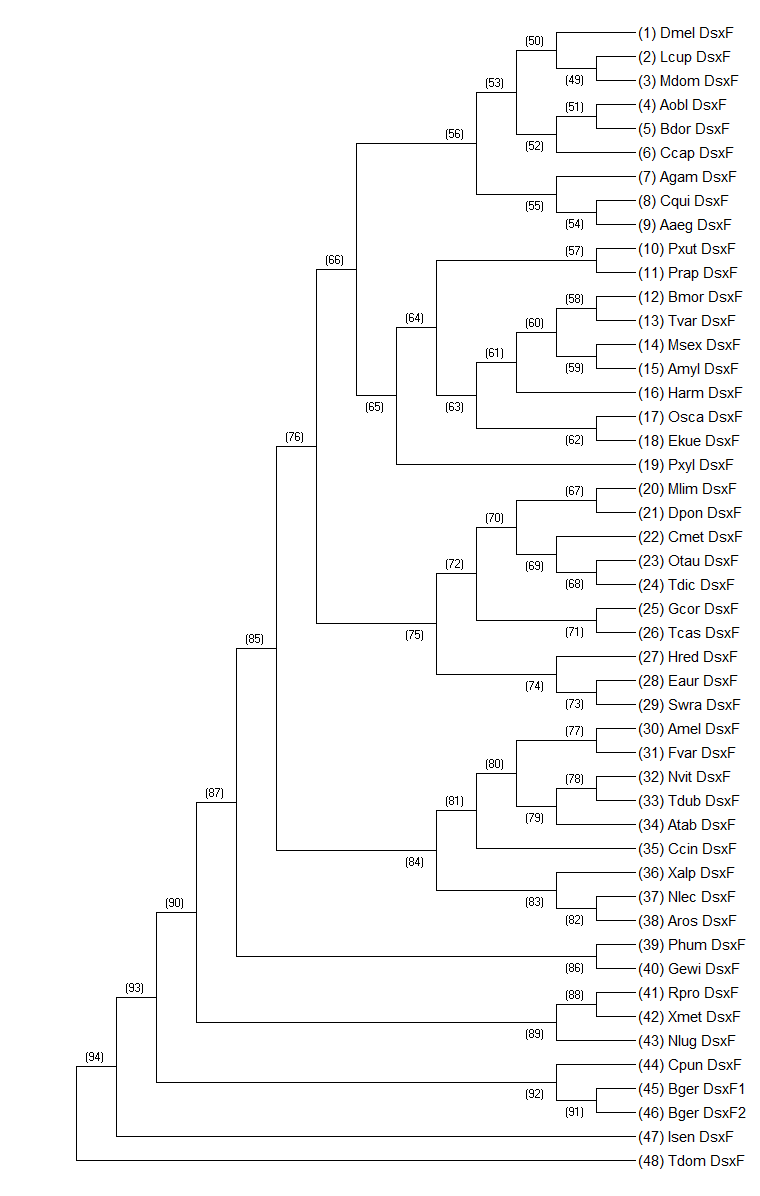
**Supplementary FIG. 9.** The guide tree used for the ancestral sequence reconstruction. The tree topology was reconstructed based on previous phylogenetic studies (Wiegmann et al. 2011; Misof et al. 2014; Li et al. 2017; Peters et al. 2017; Zhang et al. 2018; Kawahara et al. 2019; McKenna et al., 2019; Gustafson et al. 2020). The topology is here: “(((((((((((Dmel_DsxF,(Lcup_DsxF,Mdom_DsxF)),((Aobl_DsxF,Bdor_DsxF),Ccap_DsxF)),(Agam_DsxF,(Cqui_DsxF,Aaeg_DsxF))),(((Pxut_DsxF,Prap_DsxF),((((Bmor_DsxF,Tvar_DsxF),(Msex_DsxF,Amyl_DsxF)),Harm_DsxF),(Osca_DsxF,Ekue_DsxF))),Pxyl_DsxF)),((((Mlim_DsxF,Dpon_DsxF),(Cmet_DsxF,(Otau_DsxF,Tdic_DsxF))),(Gcor_DsxF,Tcas_DsxF)),(Hred_DsxF,(Eaur_DsxF,(Swra_DsxF,Ains_DsxF))))),((((Amel_DsxF,Fvar_DsxF),((Nvit_DsxF,Tdub_DsxF),Atab_DsxF)),Ccin_DsxF),(Xalp_DsxF,(Nlec_DsxF,Aros_DsxF)))),(Phum_DsxF,Gewi_DsxF)),((Rpro_DsxF,Xmet_DsxF),Nlug_DsxF)),(Cpun_DsxF,(Bger_DsxF1,Bger_DsxF2))),Isen_DsxF),Tdom_DsxF);”. The numbers on the nodes indicate the operational taxon units (OTUs) and hypothetical taxon units (HTUs) matching with the numbers of taxon names in supplementary table 8. The OTU names can be referred to in supplementary table 7.

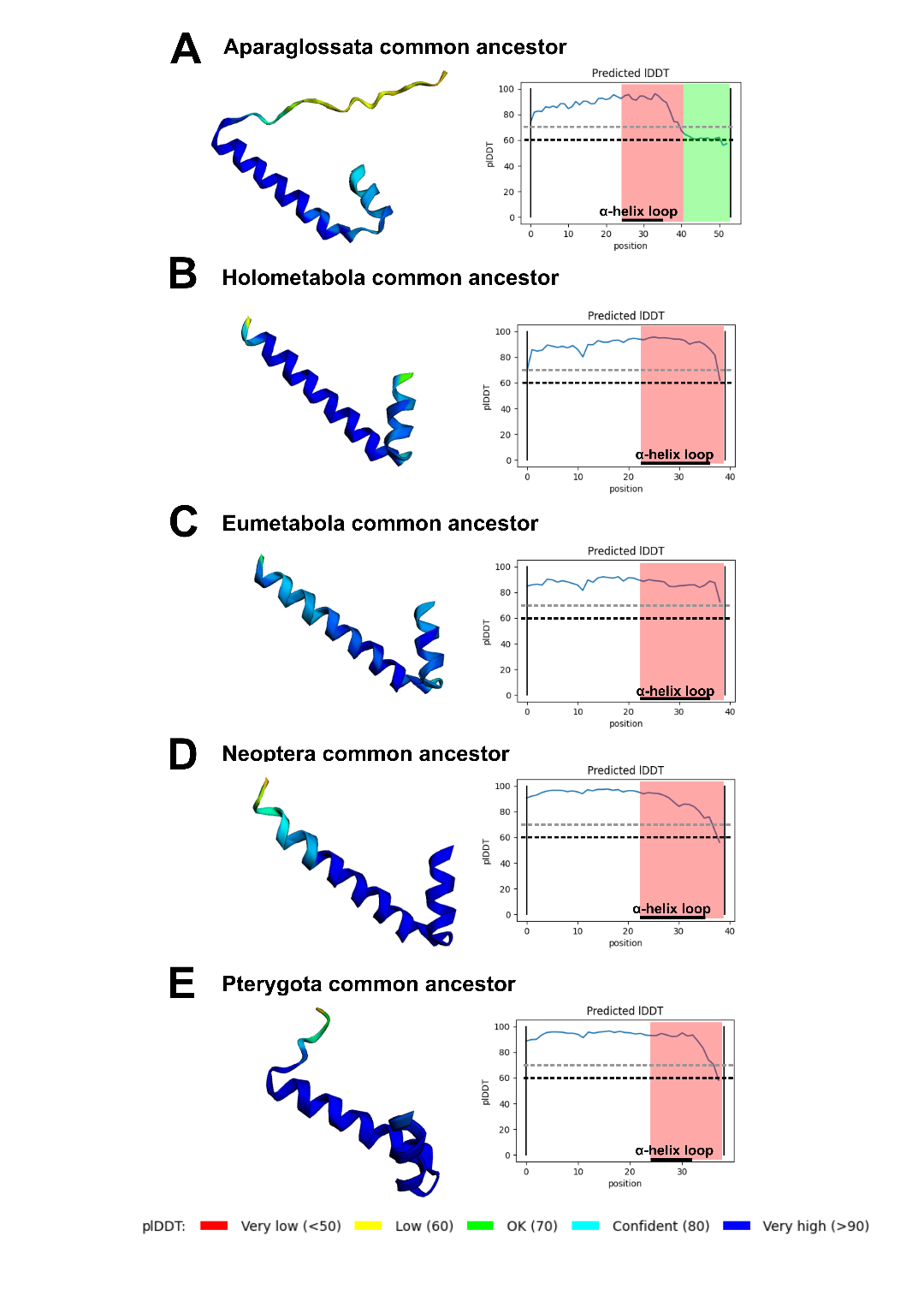
**Supplementary FIG. 10.** Accuracy of structure predictions of *dsx* female-type. Results of prediction of *dsx* female-type structure in the common ancestor of Aparaglossata (*A*), the common ancestor of Holometabola (*B*), the common ancestor of Eumetabola (*C*), the common ancestor of Neoptera (*D*), and the common ancestor of Aparaglossata (*D*). In each panel, the right 3D model shows the predicted structure of *dsx* female-type colored by its predicted local distance difference test (plDDT) score. The legend of color in the 3D model is shown at the bottom of the figure. The left graph indicates the plDDT score in each residue. The female-specific region is shown by red background. The Aparaglossata-specific region is colored by green. The black bar at the bottom of each graph shows the region predicted as an α-helix loop in the female-specific region. The black and gray dotted lines indicate plDDT = 60 and 70.

**Captions of Supplementary Tables**

**Supplementary Table 1.** Taxa and proteins used for molecular phylogenetic analysis of DMRT family.

**Supplementary Table 2.** Results of RT-qPCR assay and Brunner–Munzel test.

**Supplementary Table 3.** Results of Smirnov–Grubbs' test for expression level of *dsx* mRNA in nymphal RNAi males.

**Supplementary Table 4.** Results of generalized linear model of female traits.

**Supplementary Table 5.** Results of generalized linear model of male traits.

**Supplementary Table 6.** The expression level of *vitellogenin* genes in *Thermobia domestica*.

**Supplementary Table 7.** The taxa list used for the ancestral sequence reconstruction of *dsx*.

**Supplementary Table 8.** The result of the ancestral sequence reconstruction of the C-terminal region of *dsx* female-type.

**Supplementary Table 9.** Primers’ list used in this study.

**Supplementary Table 10.** Probabilities of reconstructed ancestral sequences of dsx female-type.
